# Analysis of Metabolomics and Transcriptomics Data to Assess Interactions in Microalgal Co-culture of *Skeletonema marinoi* and *Prymnesium parvum*

**DOI:** 10.1101/2023.12.23.573174

**Authors:** Mahnoor Zulfiqar, Anne-Susann Abel, Emanuel Barth, Kristy Syhapanha, Remington Xavier Poulin, Sassrika Nethmini Costa Warnakulasuriya Dehiwalage, Georg Pohnert, Christoph Steinbeck, Kristian Peters, Maria Sorokina

**Author notes:** Remington Xavier Poulin and Kristy Syhapanha, Department of Chemistry and Biochemistry, Center for Marine Science, College of Science and Engineering, University of North Carolina Wilmington, Wilmington, NC, USA. Anne-Susann Abel, Theoretical Biochemistry Group, Institute for Theoretical Chemistry, University of Vienna, Wien, Austria and Algorithmic Cheminformatics Group, Department of Mathematics and Computer Science, University of Southern Denmark, Odense, Denmark.

## Abstract

In marine ecosystems, microbial communities often interact using specialised metabolites, which play a central role in shaping the dynamics of the ecological networks and maintaining the balance of the ecosystem. With metabolomics and transcriptomics analyses, this study explores the interactions between two marine microalgae, *Skeletonema marinoi* and *Prymnesium parvum*, grown in mono-cultures and non-contact co-cultures. As a growth indicator, the photosynthetic potential, measured via fluorescence, suggested chemical interaction between *S. marinoi* and *P. parvum*. Using Liquid Chromatography-Mass Spectrometry (LC-MS) data, we identified 346 and 521 differentially produced features in the endo- and exometabolome of *S. marinoi* and *P. parvum*, respectively. Despite limited tandem mass spectrometry data (MS^2^) for these features, we structurally annotated 14 compounds, most of which were previously under-studied specialised metabolites. Differential gene expression analysis was then performed on the transcriptomes of the microalgae, which uncovered differentially expressed genes involved in energy metabolism and cellular repair for both species. These metabolic and transcriptomics changes depict the adaptation of both species in the co-culture. However, further data acquisition and investigation will be necessary to confirm the type of interaction and the underlying mechanisms.

**Importance:** Marine microalgae have great ecological importance and biochemical potential. Among these microbes are the diatom *Skeletonema marinoi*, known for its marine biogeochemical cycling, and the haptophyte *Prymnesium parvum*, which poses adverse environmental consequences. Given these opposing roles for the two cosmopolitan microalgae, we designed a study using untargeted metabolomics and transcriptomics to acquire a comprehensive snapshot of their interactions, grown as mono-cultures and co-cultures. The statistical analysis of the chlorophyll *a* fluorescence levels, and the metabolomics and transcriptomics dataset revealed metabolic communication occurring among the two species via specialised metabolites and activated cellular repair mechanisms. These findings reveal the complexity of the interactions within marine microbial ecosystems, offering a foundation for future research to understand and harness marine ecological systems.

## 1 Introduction

Marine ecosystems cover almost 70% of the earth’s surface, represent 95% of the known biosphere, and are a hub of undiscovered chemodiversity (1, 2). Marine microorganisms thrive in hostile environmental conditions and are constantly adapting through the exchange of genetic material and signalling metabolites (3–5). The diverse metabolites released from the interactions between microbes and their environment enhance stress tolerance and their survival capabilities (6, 7). These marine metabolites have potential applications in pharmaceuticals, biofuels, and agriculture (8). Despite these benefits, research on microbial interactions is challenging due to the complexity of microbial communities, which requires inventive experimental and *in silico* techniques that combine existing and novel methodologies to understand the metabolic exchange involved in metabolic communication (9).

Exploration of natural products (NPs) in microbial communities is conducted in various ways. Bioassay-guided fractionation is one of the classic techniques used in drug discovery to isolate and characterise bioactive NPs (10). Since the emergence of omics tools, such as genomics, transcriptomics, and metabolomics,meta-analyses of large-scale datasets have been extensively used to determine the taxonomic and metabolic profiles of marine microorganisms, enabling elucidation of biomolecules and their potential roles within the microbial communities (11). Combined transcriptomics and metabolomics data can accelerate NP discovery by analysing differential gene expression and differentially produced metabolites within the community (12, 13). However, deriving meaningful insights from such data analysis is challenging. Automating omics data analysis is essential for easier comprehension of biological processes and overcoming the computational resource limitations of managing and analysing omics data (14).

Co-cultures of two microorganisms could serve as a baseline to study complex interactions among microbes via omics techniques. For this project, we cultivated two unicellular marine microalgae, *Skeletonema marinoi* and *Prymnesium parvum*, in a mono- and co-culture experimental setup (15, 16). *Skeletonema marinoi*, a centric diatom, is known for its beneficial role in the marine food chain and biogeochemical cycling of carbon, nitrogen and silica (17, 18). *Prymnesium parvum*, also known as “golden algae”, is a marine and freshwater haptophyte that releases toxins called prymnesins and is involved in causing harmful algal blooms (HABs) (19, 20). Given the contrasting observed roles of the two ubiquitous microalgae, we designed a co-culture experiment based on the possibility of metabolic interactions among them. We implemented a workflow to analyse co-culture metabolomics data from the endo- and exometabolome of two microorganisms. Additionally, transcriptomics data were acquired under the same conditions. Statistical analyses, including Principal Component Analysis (PCA) and Partial Least Square (PLS), were performed to identify differentially produced metabolic features and differentially expressed genes. This research is a step forward in unravelling the complexities of marine microbial interactions using omics approaches.

## 2 Results

### 2.1 Chlorophyll *a* fluorescence measurement as a growth indicator

In this co-culture experiment, *Skeletonema marinoi* and *Prymnesium parvum* were cultivated in a co-culture chamber as 1) control groups represented by mono-cultures of each microalga individually, and 2) treatment groups, represented by co-culture of both species grown together, but separated from each other by a permeable membrane barrier. The chlorophyll *a* fluorescence was measured for eight days for all groups. One of the potential mechanisms of actions of allelopathy, the process of chemical exudates from one phytoplankton to affect a co-occurring competitor, is via inactivation or modulation of photosynthetic efficiency (21). However, we are working under the assumption of a linear correlation between the growth of microbial cells and the increase in chlorophyll *a* concentration. The chlorophyll *a* fluorescence over the eight days of culturing the microbial cells in mono-culture and co-culture for both species are shown in Figure 1, derived from Supplementary Table 1. Figure 1 (a) shows a lower chlorophyll *a* fluorescence in *S. marinoi* when grown as a co-culture in the presence of *P. parvum*, compared to mono-culture controls. Figure 1 (b) shows a steady increase in the chlorophyll *a* fluorescence of *P. parvum*, indicating no difference in the co-culture *P. parvum* compared with mono-culture controls, except for at day 8. Growth of all cultures throughout the experiment indicates that they were not nutrient-limited.

**Figure 1:**
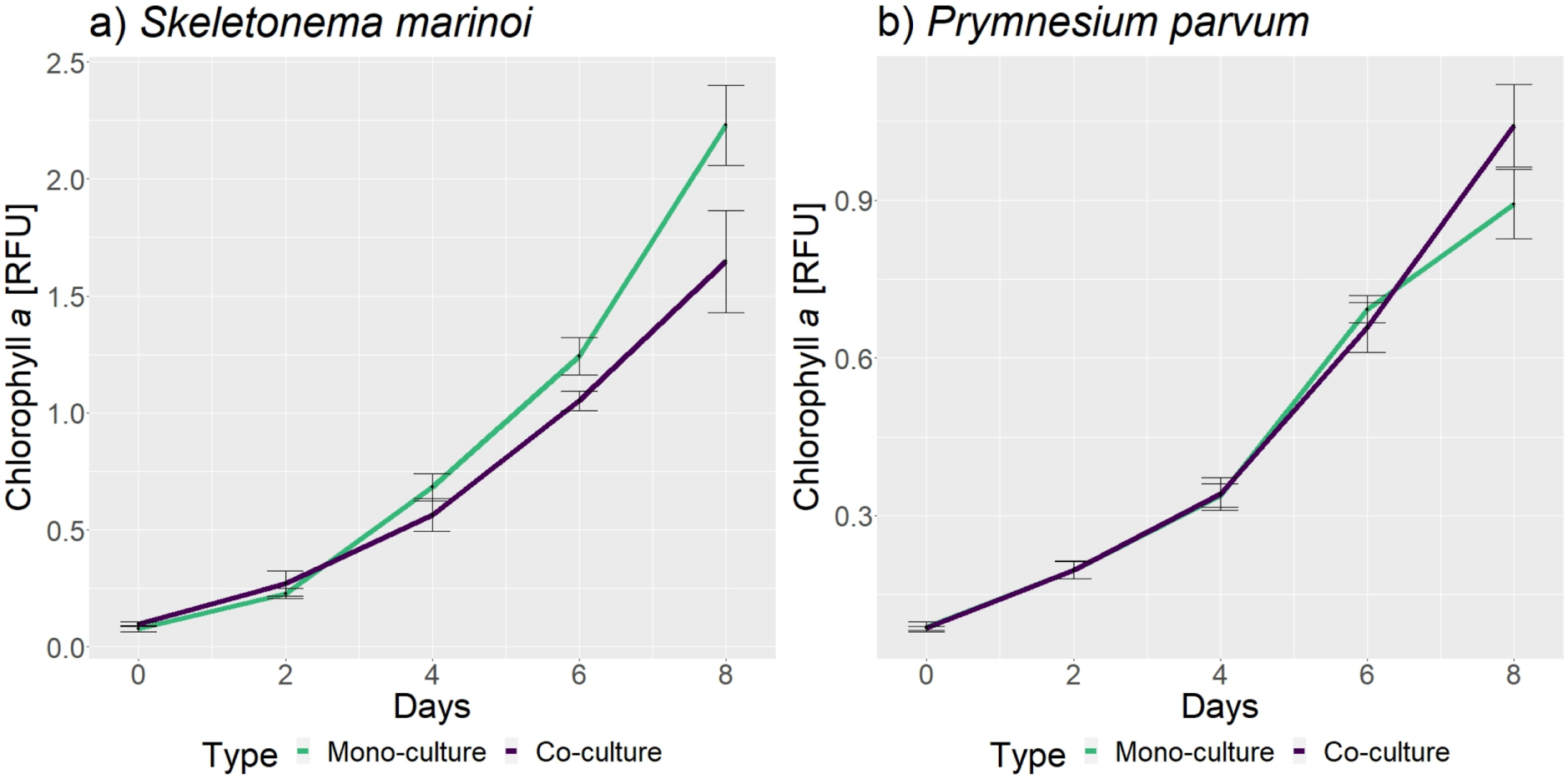
Chlorophyll *a* concentration measurement in the cells for eight days. Both plots show the number of days on the x-axis and the chlorophyll *a* fluorescence measurement in relative fluorescence units (RFU) on the y-axis. All eight replicates for monocultures and all 4 replicates for co-culture were used to generate the fluorescence plot of the cultures over eight days (see Supplementary Table 1). The error bars represent the standard deviation among the mean of the replicates for the particular day. a) The plot for the chlorophyll *a* measurement of *S. marinoi* shows lower chlorophyll *a* fluorescence of the co-culture (purple) as compared to the mono-culture (green) as a general trend. b) The plot for the chlorophyll *a* measurement of *P. parvum* shows no distinction between the co-culture (purple) and the mono-culture (green) groups. However, there is an increase in the chlorophyll *a* concentration compared to the control after day 6 in the co-culture.

Based on the chlorophyll *a* measurements plot, it can be suggested that *P. parvum* negatively influences the growth of *S. marinoi,* while *S. marinoi* doesn’t influence the growth of *P. parvum* for the first six days of growth as a co-culture. We performed a two-way ANOVA test with replicates (Table 1) to confirm this hypothesis. ANOVA was performed with 4 replicates for co-culture and 4 replicates for mono-culture for each species separately (Supplementary Table 2). For *S. marinoi*, the p-value for the variation in sample replicates over eight days is significant, while for *P. parvum* is slightly higher than the threshold of 0.05. To compare the individual days after significant p-value for fluorescence measurements of *S. marinoi* cultures in ANOVA, we performed a t-test to verify which days had significant differences in the chlorophyll *a* fluorescence. Despite non-significant findings in the ANOVA analysis of the *P. parvum* growth curve, a notable visual difference in the curves on day 8 led us to conduct a t-test on the chlorophyll *a* fluorescence in *P. parvum* cultures. Table 1 shows the t-test p-values for individual days for *S. marinoi* and *P. parvum*. The t-test performed on *S. marinoi* fluorescence data shows significant differences (p-value =< 0.05) for days 0, 4, 6, and 8. While for *P. parvum*, the p-value only dropped below 0.05 at the concentrations measured for day 8, which confirms the chlorophyll *a* measurement plots in Figure 1.

**Table 1:** Two-way ANOVA with replicates and Welch t-test (two-tailed t-test with a sample of unequal variance) to evaluate the difference in variation between mono-culture and co-culture data for *S. marinoi* and *P. parvum* on the days of sampling. To reject the null hypothesis (no variance in mono-culture and co-culture chlorophyll *a* measurements for each species is observed), the p-value is greater than 0.05, while the alternative hypothesis (significant difference between the chlorophyll *a* measurements for mono-culture and co-culture for each species) is indicated by p-values less than or equal to 0.05, which are highlighted as green in the table.

|  | ANOVA<br>p-value | day 0 | day 2 | day 4 | day 6 | day 8 |
| --- | --- | --- | --- | --- | --- | --- |
| *S. marinoi* | 5.74E-05 | 0.0232 | 0.2037 | 0.0334 | 0.0003 | 0.0058 |
| *P. parvum* | 0.0674 | 0.6711 | 0.9375 | 0.8747 | 0.2481 | 0.0214 |

### 2.2 Metabolomics analysis of endo- and exometabolome

#### 2.2.1 Statistical evaluation of differentially produced metabolites from MS^1^ data

The metabolomics MS^1^ spectra were processed to investigate observed growth effects using metabolomics preprocessing pipeline from XCMS. Overall, 24,862 features were detected for the endometabolome and 34,671 features for exometabolome from the whole dataset. After preprocessing was performed, various statistical tests were applied to the endo- and exometabolome of the co-cultures and mono-cultures. To assess the chemodiversity, Shannon diversity (H’) index was used, where higher values indicate greater diversity. In both cases, the co-culture had a higher H’ index than the mono-cultures for the same species, depicting higher chemodiversity (Supplementary Figure 1). Partial Least Square (PLS) analysis was performed to assess which spectral features caused this chemical diversity and are differentially produced among differentof *S. marinoi* and *P. parvum* (Supplementary Figure 2). The PLS analysis of the endometabolome of *S. marinoi* resulted in the identification of 22 differentially produced metabolites (i.e., differentially produced features), where 7 were also present in mono-culture conditions, while 16 were only abundant in co-culture conditions. For *P. parvum*, all 30 differentially produced metabolites were significantly more abundant in the co-culture condition. The exometabolome of *S. marinoi* resulted in 324 differentially produced metabolites, of which 304 were more abundant in co-culture conditions. Similar results were obtained for the exometabolome of *P. parvum*, where among the 491 features, 418 metabolites were more abundant in the co-culture condition.

#### 2.2.2 Metabolic exchange in endo- and exometabolome

To verify the effects observed through the Shannon diversity index and PLS plots, a comparative PCA analysis of the exometabolome was performed. The first PCA plot shown in Supplementary Figure 3a depicts all exometabolome conditions (PC1 = 24.9%, and PC2 = 7.1%). Each condition forms a separate cluster on the PCA plot. The second PCA plot shown in Supplementary Figure 3b is obtained by calculating the sum of half of the mono-culture conditions (8 mono-culture replicates for each species/ 2) to be comparable with the 4 co-culture replicates per species (PC1 = 16.7%, and PC2 = 11.6%). The combined mono-culture conditions clustered together, while the co-culture conditions formed separate clusters for *P. parvum* and *S. marinoi*. The comparison PCA plot verified that the changes in the exometabolome are not a combined effect of the individual species’ mono-culture exometabolome; instead, the observed difference is due to the actual adaptation or metabolic exchange occurring in the co-culture condition.

To analyse the metabolic exchange among the species in co-culture, we searched for shared metabolites in the endometabolome of the two microalgae in co-culture separately, as the exometabolome in the co-culture conditions is mixed and cannot be assigned to one particular species.

We searched for all 30 differentially abundant metabolites found in the co-culture endometabolome of *P. parvum* within *S. marinoi* conditions to identify shared differential metabolites. Among these 30 metabolites, only 7 were exclusive to *P. parvum* co-culture condition, suggesting that 7 of the 30 metabolites are only produced by *P. parvum* in the presence of *S. marinoi*. 10 metabolites were found in the exometabolome of mono-culture *S. marinoi*, suggesting that these metabolites were produced and released by *S. marinoi* in the extracellular environment (in both mono- and co-cultures) and when produced in co-culture, were taken up by *P. parvum*. 6 metabolites were found in both mono- and co-culture endometabolomes of *S. marinoi*, which P. parvum also takes up. The remaining 7 metabolites were shared in all the above-mentioned conditions.

A similar search was performed for the shared metabolites in the co-culture endometabolome of *S. marinoi*. In this case, out of 15 differentially abundant metabolites produced by *S. marinoi* cells (endometabolome) in co-culture conditions, 7 metabolites were found to be exclusively from *S. marinoi* co-culture source in general. 8 of these 15 metabolites were found in exometabolome of mono-culture *P. parvum*, out of which 3 were also present in both endometabolome conditions of mono- and co-culture of *P. parvum*. This suggests that 3 metabolites are produced by *P. parvum* cells regardless of *S. marinoi* presence, and in total, 8 of the released metabolites produced and released by *P. parvum* in the extracellular space as mono-culture (and also produced and released in co-culture conditions) are taken up by *S. marinoi*. The shared metabolites, along with the number of differentially produced metabolites and the annotated metabolites, are depicted in Figure 2.

**Figure 2:**
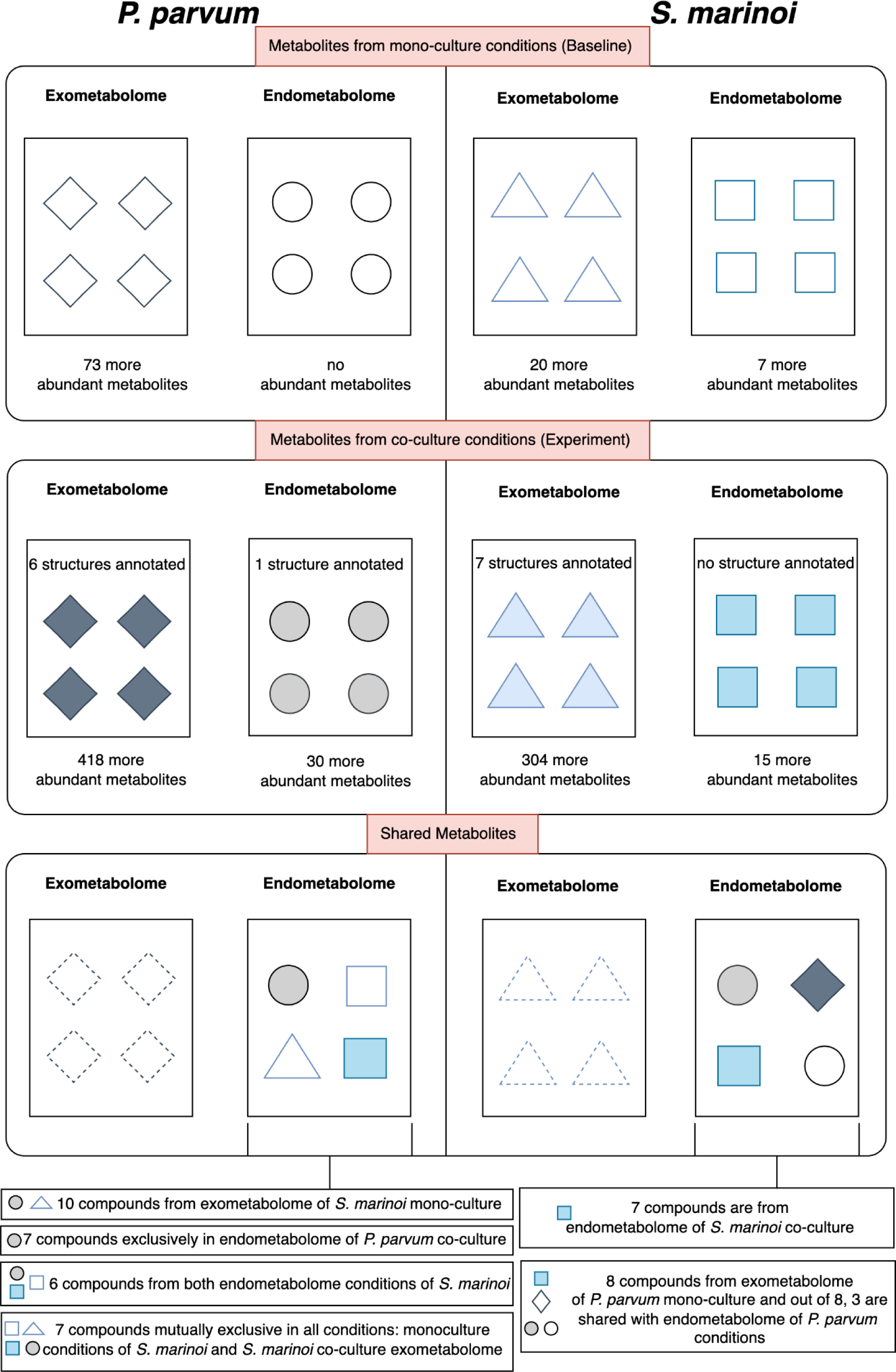
Abundant metabolites in experimental conditions. The first section refers to the mono-culture of both microalgae in endo- and exometabolome. The metabolites are depicted in each section with a coloured outline and no colour filling. The second section refers to the experimental condition of co-culture for both microalgae in endo- and exometabolome, with coloured filling representation. With our dataset, we could only annotate the metabolite structures in co-culture conditions. The third section refers to the shared metabolites which were detected in the co-culture endometabolome of both microalgae. We do not consider the shared metabolites in extracellular space, which is represented by a dotted outline. 30 more abundant metabolites were found in the co-culture endometabolome of *P. parvum* (as compared to mono-culture). 10 of these compounds were found in the mono-culture exometabolome of *S. marinoi*, 7 were exclusive to the endometabolome of *P. parvum* co-culture, 6 compounds were found in both (mono- and co-culture) endometabolomes of *S. marinoi* while remaining 7 compounds were found in all conditions. 15 more abundant metabolites were found in co-culture of *S. marinoi* endometabolome. Out of the 15 of these metabolites, 6 were exclusive to *S. marinoi* endometabolome co-culture, 8 metabolites were also present in mono-culture exometabolome of *P. parvum*; among these 8, 3 compounds were also present in both (mono- and co-culture) endometabolomes of *P. parvum*.

#### 2.2.3 Structural annotation and classification of MS^2^ data

The MS^2^ fragmentation spectra were acquired in data-dependent acquisition (DDA) mode for the precursor masses (*m*/*z*) in the inclusion list of selected high-intensity peaks from MS^1^ spectra. MAW (22) was used to annotate chemical structures from spectral databases and the COCONUT database (23) to the MS^2^ spectra. The number of features annotated within each MSI level is given in Table 2. No internal standards were used in this study (hence, no MSI level 1 structure identifications were assigned with MAW).

**Table 2:**
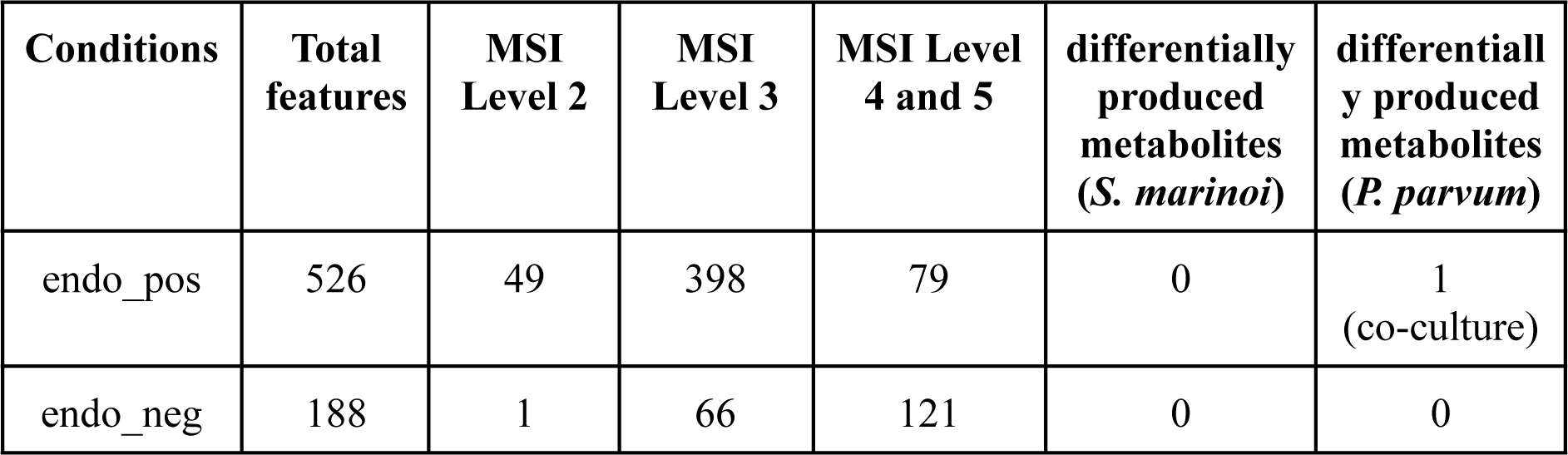
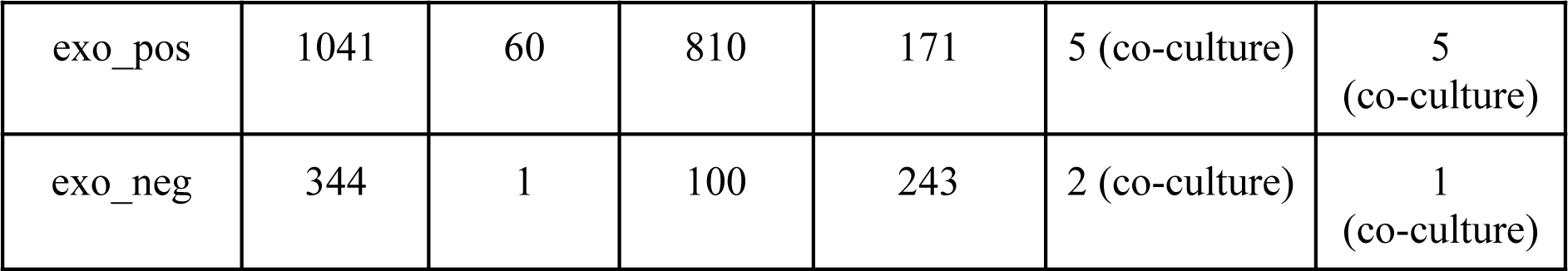
Experimental MS^2^ conditions, the associated total number of features, the features annotated together with the MSI level, and the number of differentially produced features with MS^2^ spectra. MSI level 2 refers to all features with a spectral match in a spectral database, MSI level 3 refers to all features with an annotated structure from the COCONUT database, and MSI levels 4 and 5 refer to all the rest of the features (either with chemical class, formula or mass spectrometry measurements). The conditions refer to endometabolome positive mode (endo_pos), endometabolome negative mode (endo_neg), exometabolome positive mode (exo_pos), and exometabolome negative mode (exo_neg).

With the DDA mode, only a few differentially produced metabolites were fragmented in the MS^2^ acquisition. Among the significant features of the endometabolome, the data acquired by the negative mode revealed no differentially produced metabolites, while the positive mode acquired only one in *P. parvum* co-culture condition. 10 differentially produced metabolites were annotated from exo_pos, and 3 by exo_neg samples (Table 2). The annotation results for differentially produced metabolites from MAW, represented in Table 3, unveiled the occurrence of secondary metabolites from various chemical classes.

**Table 3:**
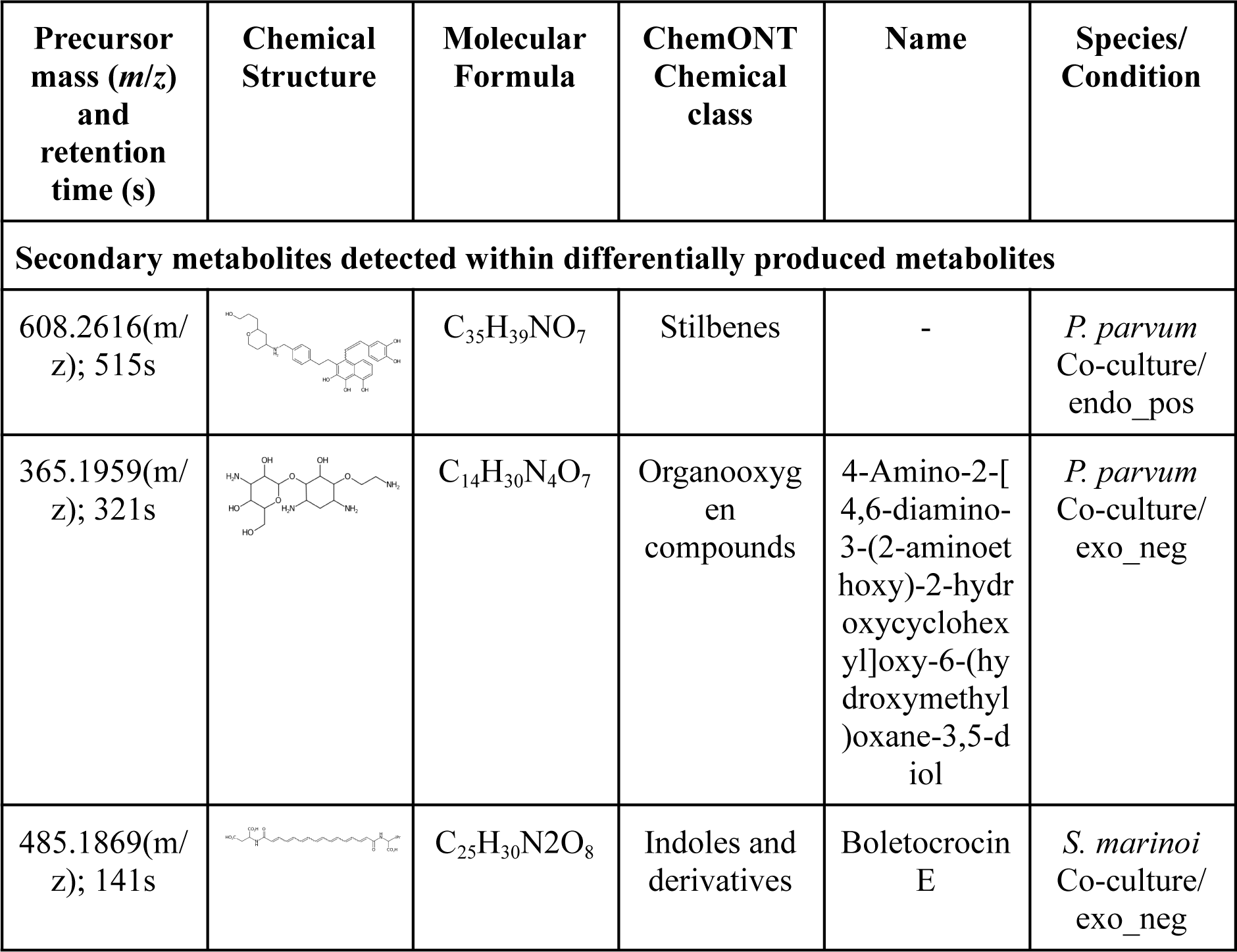
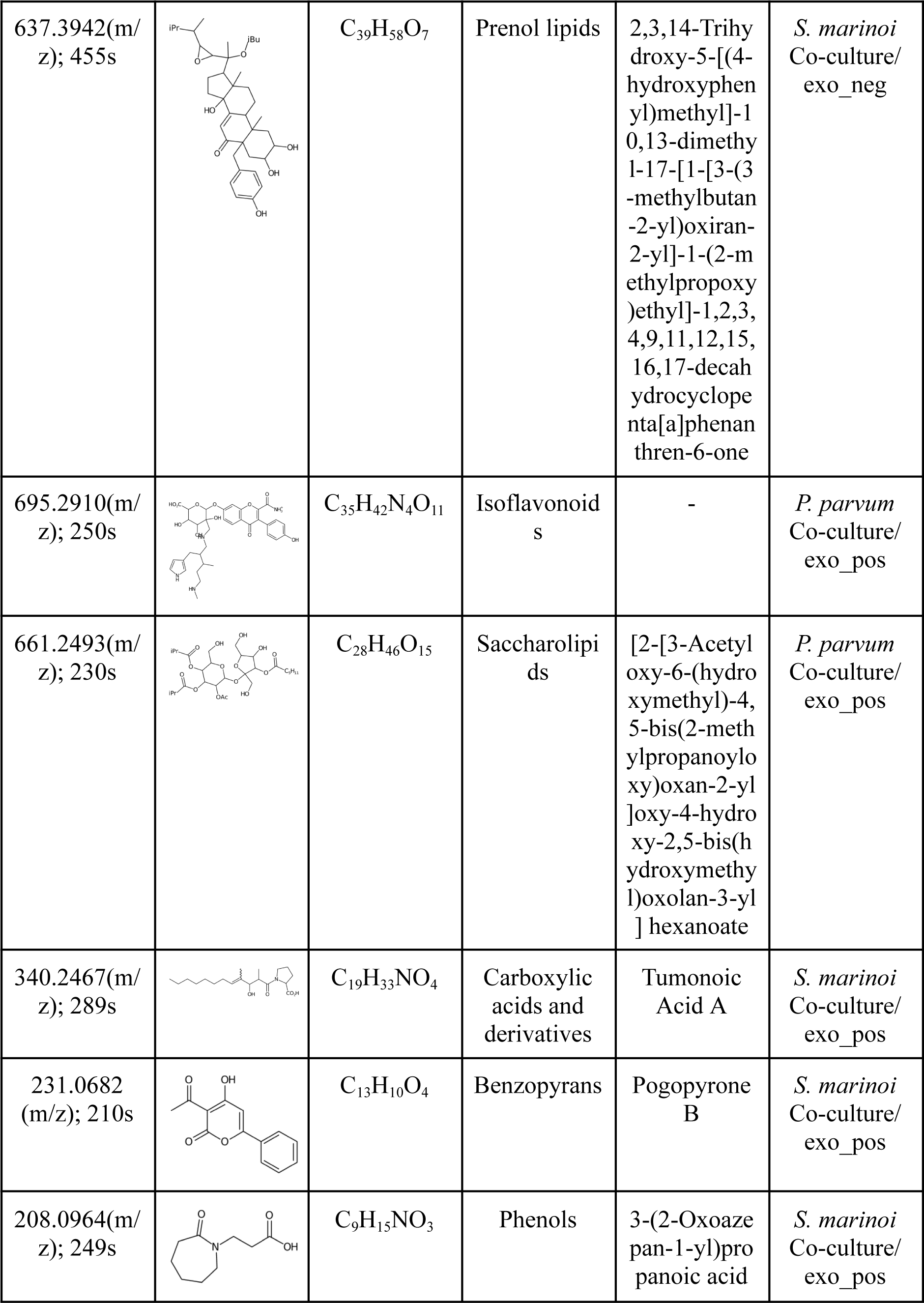
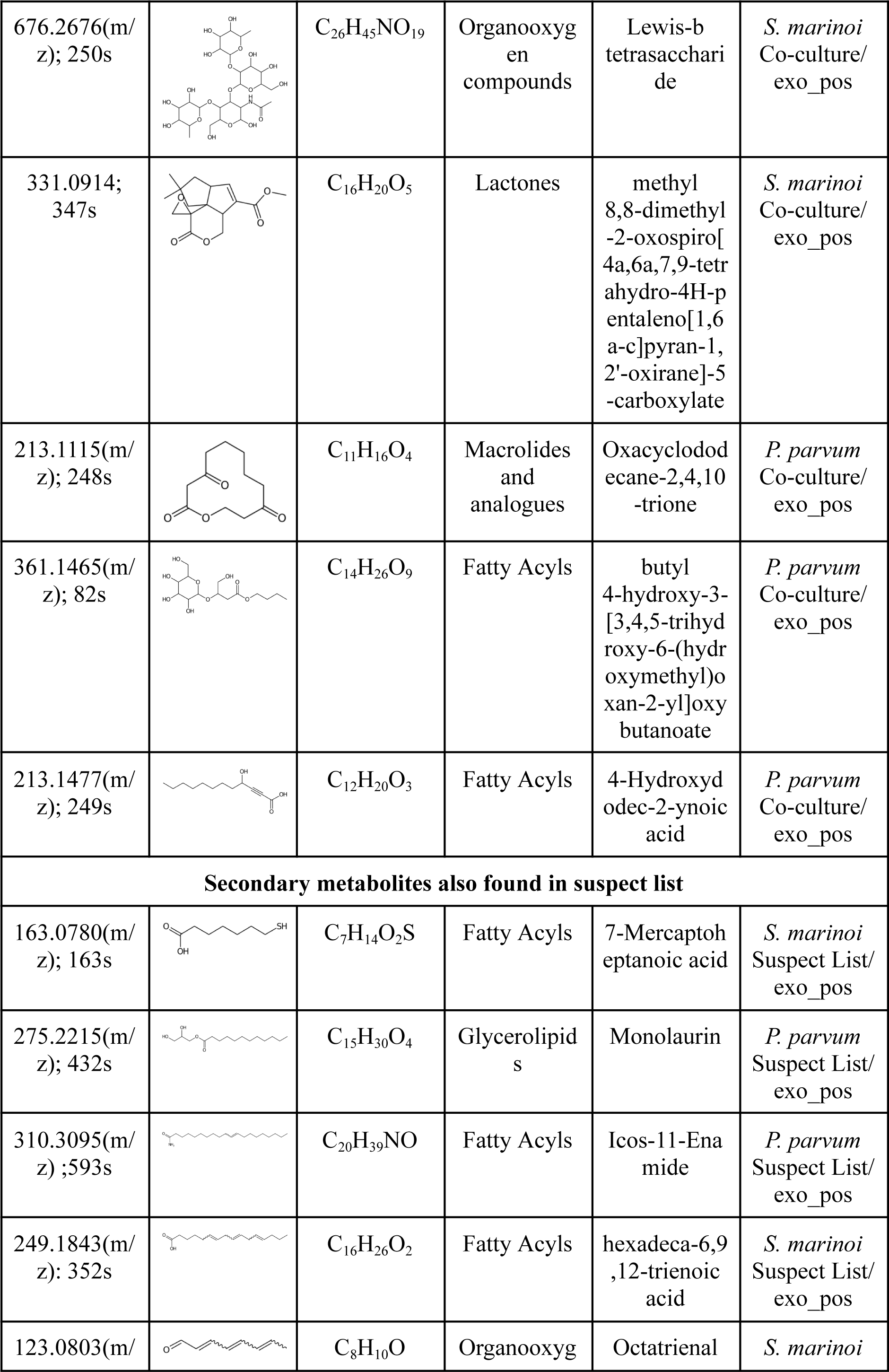
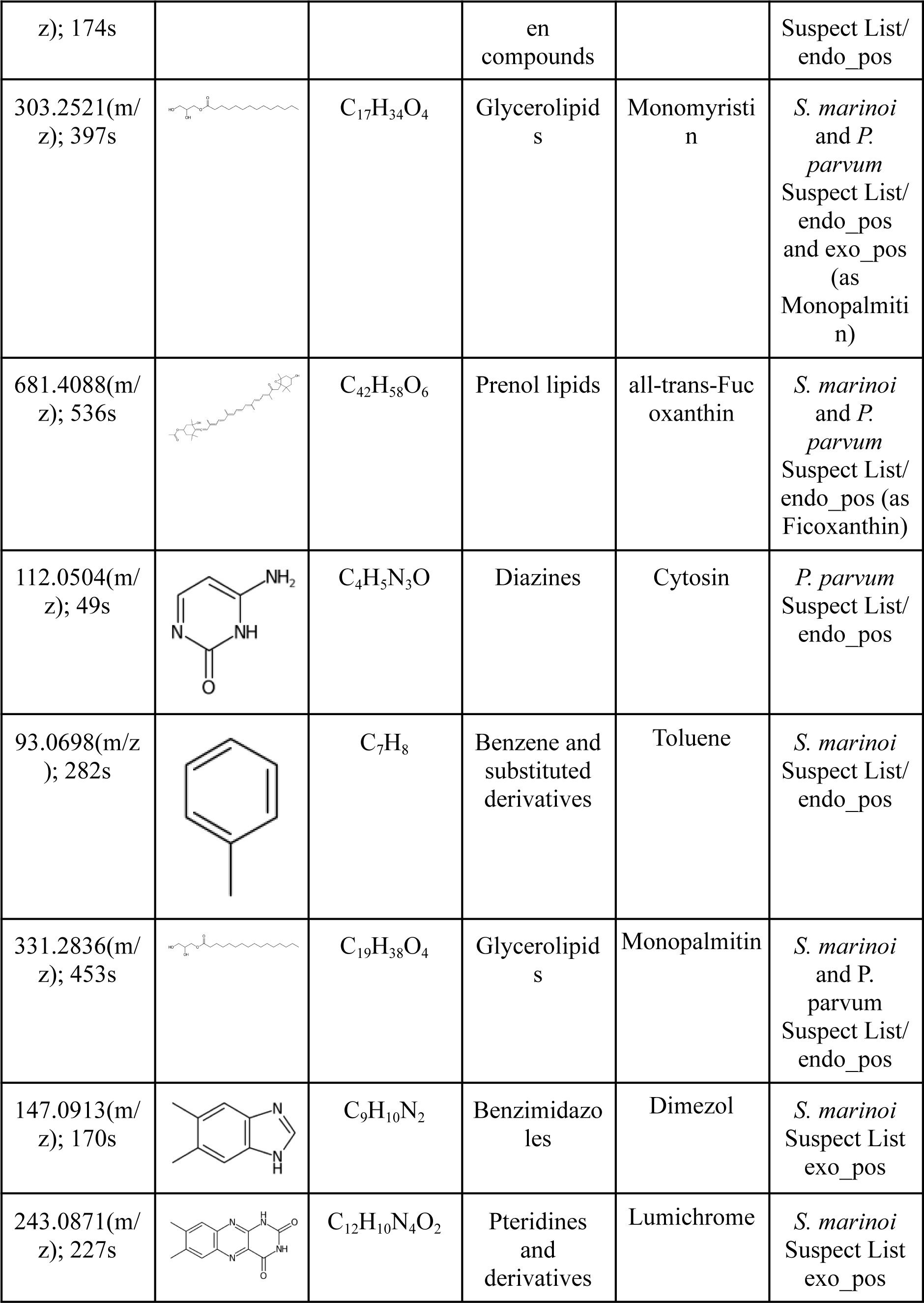
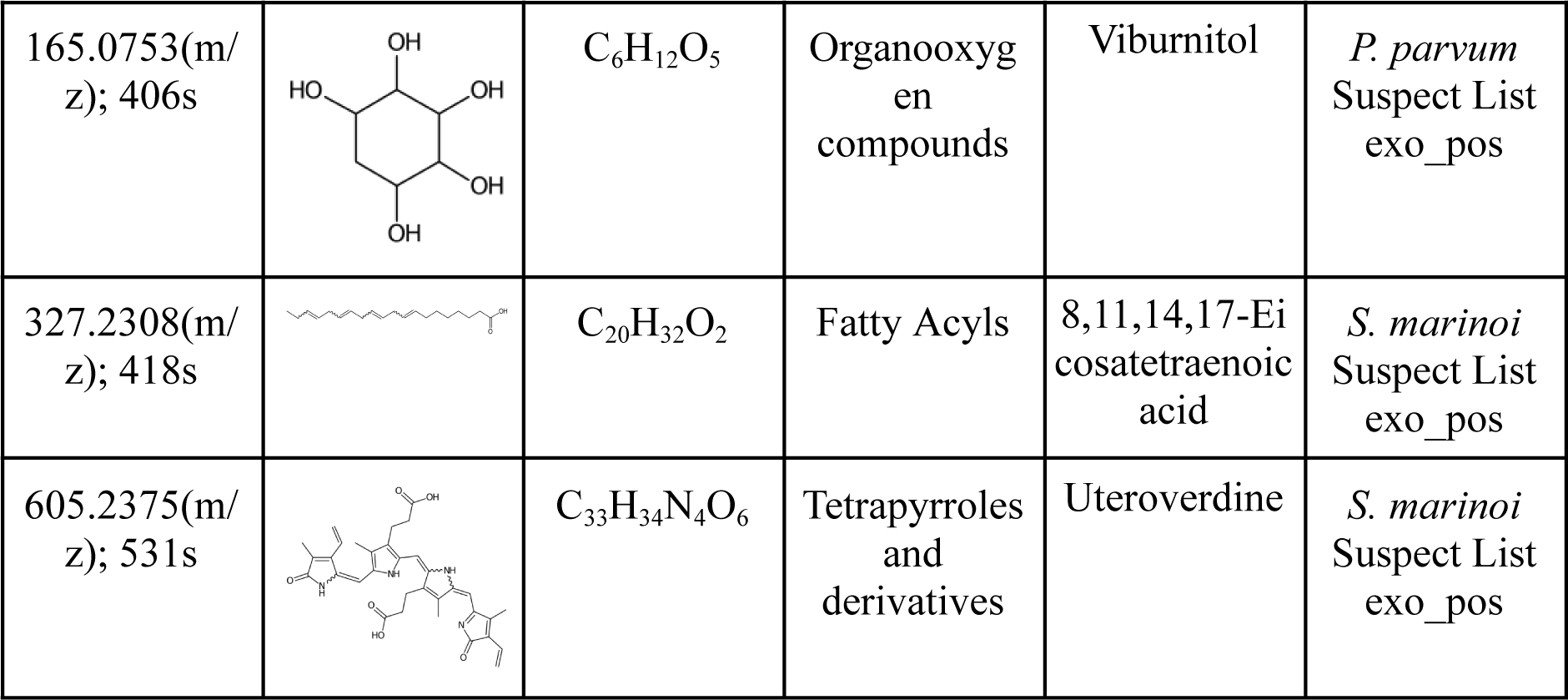
Chemical structure annotation of significant features with MAW. The precursor mass [*m*/*z*], median retention time in seconds, molecular formula, molecular structures depicted with CDK Depict (24, 25), chemical class based on ChemONT, and lastly, the origin of the differentially produced metabolites are presented in the table.

### 2.3 Transcriptomics analysis to assess differentially expressed genes across mono-culture and co-culture

Using transcriptomics data, we investigated the dynamics of gene expression in mono-cultures and co-cultures. The transcriptome assemblies for *S. marinoi* and *P. parvum* were performed with the Trinity tool (26). Within the assembly for *S. marinoi*, kraken2 (27) predicted 25% of the reads to be from humans, 24% from bacteria, 38% with no hits, and ∼1% from viruses and other sequences. About 12% of assembled transcripts were assigned directly to *S. costatum* or the group of Stramenopiles. After the quality check, Busco analysis (28) was performed to measure the quality of the transcriptome: non-restrictive transcriptome, which had both transcripts that belonged to *S. marinoi* and unclassified transcripts, and restrictive transcriptome, which had transcripts that only belonged to *S. marinoi*. Since the restrictive transcriptome had a high score of 96%, we went further with the restrictive transcriptome assembly. For *P. parvum*, the measured contamination led to bacterial (32%) and human (17%) contamination within the sequenced samples and from virus and other sequences (∼0.5%). Approximately 3% of assembled transcripts were assigned to *S. costatum* (more precisely to Stramenopiles), which are most likely *P. parvum* transcripts closely related to the Stramenopiles group, which was the closest one to *P. parvum*. Most *P. parvum* transcripts (47%) were within the unclassified group. So, the non-restrictive transcriptome, with a score of 80.7%, was used for further analysis.

Using the tool Augustus (29), 46598 of the assembled transcripts were annotated as potential Open Reading Frames (ORFs) for *S. marinoi,* and 123266 were annotated for *P. parvum*. From within these transcripts, Augutus classified 32478 ORFs for *S. marinoi* and 59328 for *P. parvum* with functional domains. For *S. marinoi*, Augustus predicted that the assembled potential transcripts could encode for more than one ORF. For many of these transcripts, the predicted coding sequences (CDS) are located on opposite strands, i.e., one CDS was predicted in the sense direction of the assembled transcript and the other CDS in the anti-sense direction. To better estimate the number of unique transcripts, MMSeqs2 (30) was used. It assembled the transcripts based on the proteins they encoded, resulting in 10812 transcripts for *S. marinoi* and 75575 for *P. parvum* (Table 4).

**Table 4:** Descriptive Statistics for *S. marinoi* and *P. parvum* transcriptome assembly.

| Species | Assembled potential transcript sequences by Trinity | Total Annotated Potential ORFs by Augustus | Classified ORFs by Augustus | Total unique ORFs |
| --- | --- | --- | --- | --- |
| *S. marinoi*<br>(restrictive transcriptome) | 31958 | 46598 | 32478 | 10812 |
| *P. parvum*<br>(non-restrictive transcriptome) | 230604 | 123266 | 59328 | 75575 |

The transcriptome assembly with ORFs predictions was then followed by the differential gene expression analysis to identify the differentially expressed ORFs (Figure 3). The differentially expressed ORFs with an FDR-adjusted p-value less than 0.05 and sorted genes based on the log fold change values resulted in 664 differentially expressed ORFs for *S. marinoi* and 755 for *P. parvum*. Table 5 lists the top 10 differentially expressed ORFs from *S. marinoi* and *P. parvum* with their functional annotation when identified. The transcriptome sequences are available with the BioProject ID PRJNA1004186, while the differential ORF protein sequences are available on Zenodo with the DOI: 10.5281/zenodo.10397384 (31).

**Figure 3:**
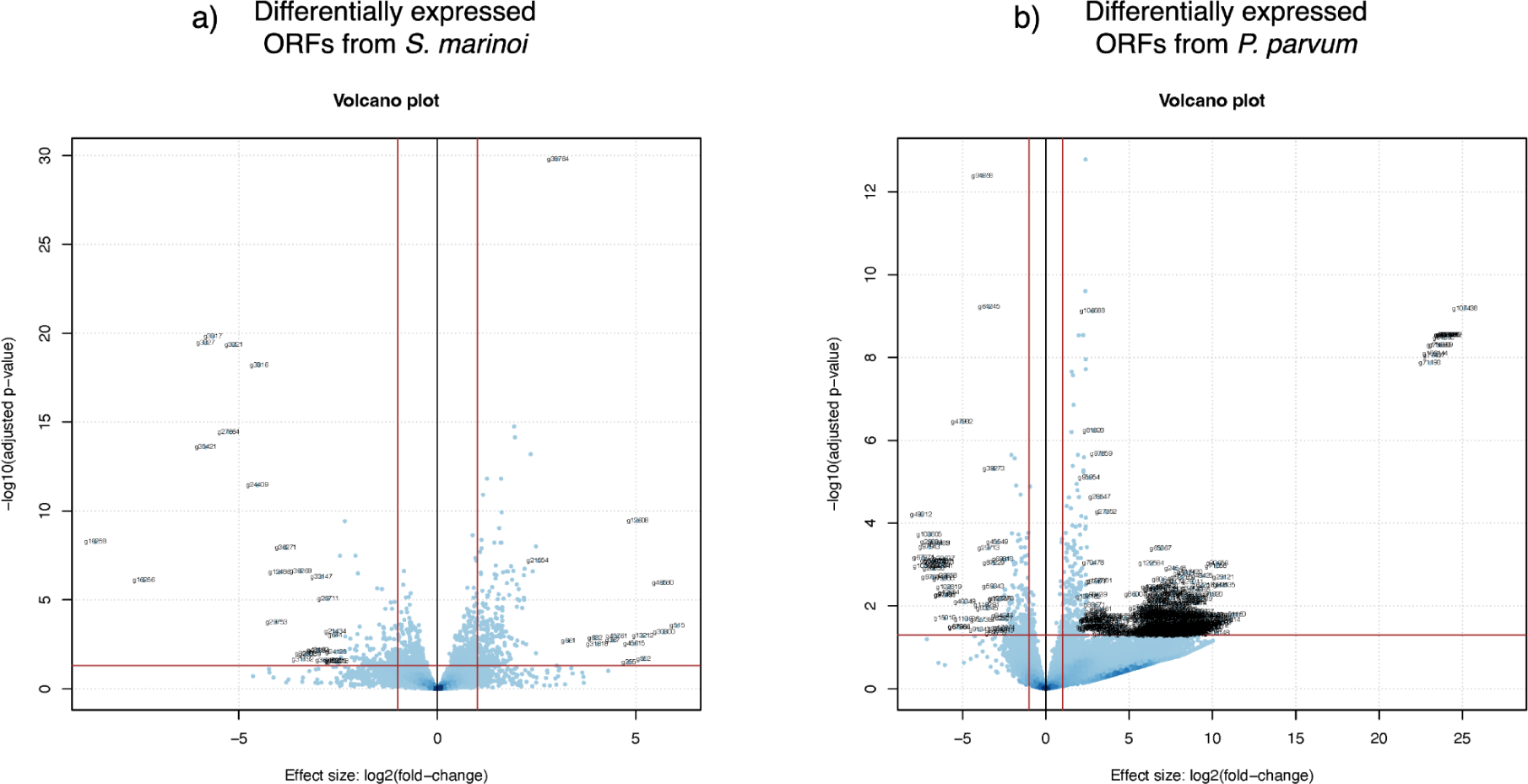
Volcano plots for the differentially repressed genes. This scatter plot displays the relationship between the log fold change and the adjusted p-value of each gene (represented by the dots). The tentative names are also displayed for the top 30 differentially expressed ORFs. The horizontal line marks the adjusted p-value threshold, whereas the two vertical lines indicate possible log fold change thresholds (1 and -1).

**Table 5:** Top 10 differentially expressed ORFs from S. marinoi and P. parvum, with putative gene ID, log2 fold change, where high positive values indicate its differential abundance in the co-culture and absence in the mono-culture, padj as the FDR-adjusted p-value and lastly the predicted function or domain of the differentially expressed ORFs.

| Local ORF ID | log2 fold change | Predicted domain | Predicted function |
| --- | --- | --- | --- |
| **Differentially expressed ORFs from *S. marinoi*** |  |  |  |
| g515 | 6.0440 | 26s proteasome non-atpase regulatory subunit; jab1/mov34/mpn/pad-1 ubiquitin protease; eukaryotic translation initiation factor 3 -related; maintenance of mitochondrial structure and function | ubiquitin-dependent protein catabolic process; mitochondrion organization |
| g30300 | 5.7082 | beta-ketoacyl-[acyl-carrier-protein] synthase iii, c/n-terminal; 3-oxoacyl-[acyl-carrier-protein] synthase | fatty acid biosynthetic process |
| g46580 | 5.6788 | flavin containing amine oxidoreductase; protoporphyrinogen oxidase | oxidative deamination of primary and secondary amines; porphyrin-containing compound biosynthetic process |
| g552 | 5.1935 | anaphase-promoting complex subunit 4 wd40 domain; the wd repeat cdc20/fizzy family; fizzy-related 2 isoform 1; anaphase-promoting complex subunit 4 | anaphase-promoting complex-dependent catabolic process |
| g13212 | 5.1754 | secy/sec61-alpha family | protein translocation channel |
| g12608 | 5.0477 | unknown domain | unknown function |
| g45615 | 4.9562 | enolase | glycolysis |
| g255 | 4.8223 | unknown domain | unknown function |
| g45761 | 4.5164 | acyl-coa desaturase | unsaturated fatty acid biosynthetic process |
| g367 | 4.4115 | small gtpase rho/ras family profile | small GTPase mediated signal transduction |
| **Differentially expressed ORFs from *P. parvum*** |  |  |  |
| g107438 | 25.1531 | putative zinc-binding metallo-peptidase; adp-ribosylglycohydrolase; | Proteolysis; protein de-ADP-ribosylation |
| g87732 | 24.3294 | unknown domain | unknown function |
| g90799 | 24.2173 | unknown domain | unknown function |
| g68158 | 24.0612 | unknown domain | unknown function |
| g31038 | 24.0559 | major facilitator superfamily (mfs) profile | transmembrane transporter activity |
| g80644 | 24.0392 | trifunctional enzyme subunit alpha;enoyl-coa hydratase/isomerase | fatty acid metabolism |
| g64217 | 23.9817 | Transcriptional regulator MraZ; SpoVT-AbrB domain profile | division/cell wall cluster transcriptional repressor |
| g89853 | 23.9741 | macB-like periplasmic core domain | regulation of xenobiotic detoxification by transmembrane export across the plasma membrane |
| g51890 | 23.8731 | copC domain | copper transport protein |
| g114069 | 23.6307 | unknown domain | unknown function |

The top 10 differentially expressed ORFs in both species that could not be annotated with a function or domain were then investigated for an attempt of manual functional annotation, using TMHMM (32), p-BLAST (33), and InterProScan (34) for hints to identify the category of their possible function (Table 5). Among the ORFs of unknown function in *S. marinoi*, the ORF number g12608 could be predicted to encode a transmembrane protein, exhibiting 78% homology with a hypothetical protein from the haptophyte phytoplankton *Chrysochromulina tobinii*. In contrast, the ORF g45761 is likely to encode an intracellular peptide with no detectable homology. In total, 229 ORFs in the differential expression analysis in *S. marinoi* had completely unknown functions and were only homologs with genes of unknown function in public databases. For *P. parvum*, four ORFs among the top 10 differentially expressed ones were with unknown functions. Among the four ORFs, ORF number g87732 was predicted to encode an intracellular protein, showing no significant similarity to any known or hypothetical proteins, while ORF number g90799 was predicted to encode a transmembrane protein with otherwise no known similarity to anything else in public databases. Furthermore, the subsequent ORFs numbers g68158 and g114069 also appeared to be intracellular. The response of *P. parvum* (highest fold change is 25.1531) is way higher than *S. marinoi* (highest fold change is 6.0440). These findings highlight the considerable number of genes with yet-to-be-characterised functions in both *S. marinoi* and *P. parvum* and an increased level of differential expression in the genome of *P. parvum* when grown as co-culture with *S. marinoi*.

## 3 Discussion

### 3.1 Influence of co-culturing on microalgae

Microbial communities are complex adaptive biosystems and are highly dynamic, with varying biodiversity and chemodiversity depending on the exogenous stimuli (35). However, to understand the complexities of such systems, the controlled cultivation of two microorganisms can be considered as a basis for studying highly complex communities. Both microalgae selected for this research are cosmopolitan, found in marine ecosystems, and hold profound impacts on the aquatic biosphere (36, 37). The first microalgae in this experiment is the diatom *S. marinoi*, which is a valuable source of different natural products such as polyunsaturated fatty acids [81], serves as a food source for other marine organisms and has been demonstrated to convert CO_2_ in cement waste gas into renewable energy (39), depicting ecological and economic applications. The second microalgae, *Prymnesium parvum*, is considered a toxic species, which causes a decrease in biodiversity during harmful algal blooms (HABs) while growing at an exponential rate (40), indicating a toxic or growth-inhibiting behaviour on other members of the microbial community. It remains unexplored whether the growth of *P. parvum* benefits from diatoms like *S. marinoi* and other microalgae.

Our investigations into the co-culture of *S. marinoi* and *P. parvum* revealed distinct changes in chlorophyll *a* fluorescence, which is used as a proxy for cell growth in this experiment. Specifically, *S. marinoi* demonstrated a significant reduction in chlorophyll *a* fluorescence when co-cultured with *P. parvum* as compared to when grown as a mono-culture, whereas *P. parvum* in co-cultures demonstrated no significant difference overall, except the last day of fluorescence measurement in the cultures (as demonstrated in Figure 1 and Table 1). These observations could be due to several reasons. *P. parvum* is also known to be a mixotroph; hence, it relies on both photosynthesis and other organisms for food sources (41). The cell lytic activity of the metabolites released from *P. parvum* could cause *S. marinoi* cellular depletion and the transfer of nutrients from *S. marinoi*. Supervised statistical analysis was performed to elucidate the possible metabolic exchange via metabolomics and transcriptomics data.

### 3.2 Exploring responses to mutual presence of microalgae through metabolome and transcriptome

To investigate the metabolic interactions observed in co-culture of the two microalgae, suggested by a shift in the chlorophyll *a* concentrations, we employed the Partial Least Squares (PLS) analysis on endo- and exometabolome of all conditions from both microalgae.

This supervised statistical regression model linked the intensity of differentially produced metabolic features to their respective occurrences in the co-culture and mono-culture conditions (Supplementary Figure 2). The 22 co-culture-specific features for the endometabolome of *S. marinoi* speculate the presence of metabolites involved in the defence activation of *S. marinoi* in the presence of *P. parvum*. For *P. parvum* endometabolome 30 differentially produced metabolites were revealed, suggesting the production of *P. parvum* allelochemicals, with adverse effects on *S. marinoi*. Alternatively, these features could have also originated from *S. marinoi*, indicating nutrient uptake from *P. parvum*.

To further investigate the potential metabolic exchange and uptake between the endometabolomes of *S. marinoi* and *P. parvum*, we analysed whether the differentially produced metabolites are shared among the endometabolomes (Figure 2). First, the differentially abundant metabolites in the endometabolome of co-culture *P. parvum* were cross-referenced with the exometabolome of the mono-culture of *S. marinoi*, and both the mono- and co-culture endometabolome of *S. marinoi*. The 23 common metabolites between endometabolome (co-culture) *P. parvum* and different conditions of *S. marinoi* metabolome suggest that *P. parvum* consumes these *S. marinoi* metabolites. Furthermore, the differentially abundant metabolites in endometabolome of co-culture *S. marinoi* were cross-referenced with exometabolome of the mono-culture of *P. parvum*, and both the mono- and co-culture endometabolome of *S. marinoi*. In this scenario, atleast 8 metabolites in *S. marinoi* endometabolome in co-culture condition were from *P. parvum*; however, these metabolites can be termed as common metabolites between two species, as they were from the set of 15 metabolites found in the co-culture condition of *S. marinoi*. Either these could be the exudates exchanged between the two species, or the production of these metabolites is induced by the presence of other species. However, these speculations need further experimental confirmation.

For exometabolome (Supplementary Figure 2), we observed an increase in differentially produced metabolites as compared to endometabolome, which could be ascribed to the experimental design, given the potential for metabolic exchange across the permeable membrane. It is plausible that these features, originating from either microalga, indicate a predominant metabolic uptake direction favouring *P. parvum*. An additional factor to consider is the non-axenic nature of the mono-cultures and co-cultures. Some of these significant metabolites might be sourced from bacteria existing symbiotically within the phycosphere of these microalgal cells. While the co-culture’s exometabolome cannot be easily attributed to a specific microalga, its distinct profile suggests metabolic interaction between *S. marinoi* and *P. parvum* rather than a mere aggregation of their individual metabolomes (Supplementary Figure 3). Even with these exometabolome features, we are missing non-polar metabolites, as the data was acquired in the Reverse Phase.

Furthermore, a comprehensive transcriptome assembly coupled with protein annotations was performed for differential gene expression analysis. The resulting volcano plots (Figure 3) depict a change in responses to mutual presence by *S. marinoi* and *P. parvum* in co-culture samples; the differential gene expression analysis revealed a total of 664 differentially expressed transcripts for *S. marinoi* and 755 for *P. parvum*. The increased number of transcripts for *P. parvum* could indicate the genetic exchange from *S. marinoi* cell lysis towards the *P. parvum* side of the co-culture chamber through the permeable membrane. This has not been confirmed with the current methodologies applied to this study. Moreover, within the top 10 differentially expressed transcripts, *P. parvum* had a higher fold change for transcripts of unknown function than the log fold change of the top 10 differentially expressed transcripts in *S. marinoi* with unknown function (Table 5). These results suggest that both species undergo significant transcriptional remodelling, potentially reflecting adaptive mechanisms or competitive behaviours to thrive in the presence of the other, especially in the case of *P. parvum*.

### 3.3 Metabolomics and transcriptomics annotation analysis and technical aspects

The differential metabolite profiling from the co-culture revealed intriguing dynamics that suggest the underlying responses and adaptations of these microorganisms in a co-culture setting; however, these metabolites were not studied in the context of marine microalgae before. In *P. parvum* co-culture (endometabolome), stilbenes, a phenolic allelochemical compound class, was annotated (42), known to have roles in defence mechanisms and regulate critical signalling pathways such as JAK-STAT, MAPK, and NF-κB pathways by diminishing the transcription of inflammatory factors and thus preserving a homeostatic environment (43). Meanwhile, the exometabolome of *P. parvum* predominantly showed fatty acyls, macrolides (44), Saccharolipids (45), and Isoflavonoids, involved in protection against oxidative stress and immunomodulation (46). These detected metabolites show a survival response from *P. parvum*; however, they could also belong to *S. marinoi* as exometabolites are exchanged through the permeable membrane. *S. marinoi*, in contrast, demonstrated the presence of lactones, phenols, and prenol lipids. Certain other known compounds, such as the fungal pigment compound boletocrocin E (47), tumonoic acid A (exhibiting anti-inflammatory properties) (48), commonly found in bacteria and marine cyanobacterium *Blennothrix cantharidosmum* (49), and pogopyrone B, a benzopyran, sourced from the plant *Pogostemon heynianus*, were detected (50). These metabolites are less studied for their functions and could be due to misannotation, because of the limited chemical structures associated with marine microalgae in the databases (51). However, their putative presence suggests that as a co-culture, *S. marinoi* and *P. parvum* produce a competitive environment. We also used suspect lists from both *S. marinoi* and *P. parvum* for all MS^2^ features to identify secondary metabolites specific to either species or both, none of which were differentially produced (Table 3). Using the suspect list, the origin of specific metabolites could be identified, most of which belonged to *S. marinoi,* which is more extensively studied along with its closely related species *S. costatum*; however, *P. parvum* is mostly studied for its toxins and hence had a much smaller suspect list of known compounds and less secondary metabolite annotations from the suspect list (details in Supplementary Results 1).

MS^2^ for only 14 differential metabolites (Table 3) is likely due to the data acquisition from the inclusion lists generated using Compound Discoverer software, which focused on the most abundant MS^1^ features in the DDA mode. Since all of the co-culture analysis was performed using open source software and R packages after both MS^1^ and subsequent MS^2^ data acquisition, no prior processing or statistical analysis was performed using Compound Discoverer to pinpoint the most relevant differential features before MS^2^ data acquisition, and there’s a high likelihood that several significant features were overlooked for MS^2^ fragmentation because they were differentially abundant among different conditions and not across all samples (52). A future follow-up using open source code would be to first acquire MS^1^, statistically analyse the differentially abundant metabolites, enrich those features (as they might be lost due to low intensity throughout the samples) and acquire MS^2^ for those metabolites.

The transcriptional landscape of *S. marinoi* and *P. parvum* in the co-culture reflected both primary and secondary metabolic adaptations, as observed from the top 30 differentially expressed ORFs (only 10 differentially expressed ORFs mentioned in Table 5; the rest can be found in Zenodo (31)) for *S. marinoi* and *P. parvum*. In *S. marinoi*, the presence of genes associated with core cellular processes like DNA repair, cell signalling, apoptosis, and immune response, based on the identification of a protease related to ATP-dependent degradation of ubiquitinated proteins (53), provided the metabolic readjustments that might be necessary for survival in a shared environment. Fatty acid synthesis through the condensation reaction and enzymes associated with oxidative deamination suggest potential lipid-based defence strategies or energy storage responses as a buffer against potential resource competition in the presence of *P. parvum* (54). The genes involved in photosynthesis, such as photosystem II d2 protein, signify a possible photosynthetic activity as seen in chlorophyll *a* levels or adaptation to changes in light quality or intensity that may arise from the growth dynamics of the co-culture (55). In *P. parvum*, the identification of genes, such as ADP-ribosylation/crystallin j1, highlights possible cellular regulatory and repair mechanisms. Similarly, the enoyl-CoA hydratase/isomerase and the dihydrodipicolinate synthetase families, both central to fundamental metabolic pathways, might indicate energy utilisation. Various transporters like carboxylic acid and copper transport protein suggest an exchange of metabolites and the metabolic content in the extracellular space. Moreover, the presence of genes associated with type IV pilus inner membrane component and the division/cell wall cluster transcriptional repressor could indicate a heightened focus on biofilm formation (56).

For the transcriptome assembly of both species, we encountered a substantial number of potentially assembled transcripts and annotated ORFs mentioned in Table 4. Most likely, many of the assembled transcripts have duplicates (sequence similarity). This could result from paralogs, small assembly mistakes, and alternative splicing events. To analyse the differentials ORFs, we used the total number of annotated ORFs so as not to lose any important differential ORFs with no functional annotation. Transcriptomics data analysis provides some insights into the co-culture dynamic; however, there are many missing annotations specific to secondary metabolism, specifically for *P. parvum*. When analysed using BLAST-P, several differentially expressed ORFs with unknown functions in this co-culture experiment showed minimal homology to hypothetical proteins. Without well-defined gene sets for these species, identifying unique genes and pathways specific to the microalgae can be challenging due to limited existing data to reference. Specifically for *P. parvum*, most of the top differentially expressed ORFs suggested primary metabolism-relevant proteins, and others with unknown functions and low structural homology to known proteins could have more significant insights into the secondary metabolism of the co-culture of these two microalgae.

### 3.4 Conclusion

Chemical interactions regulate many marine life processes, such as balancing the ecosystem and the type of interaction among the microbial key players, such as predation or defence (57). Natural products which cause marine chemical interaction affect not only the endometabolome of the species but also the exometabolome in the environment. This study describes the co-culture setup between two microalgae, *Skeletonema marinoi* and *Prymnesium parvum*. The cell growth data suggested a negative growth impact on *S. marinoi* when grown in the presence of *P. parvum*. This hypothesis was then tested with statistical analysis, metabolomics, and transcriptomics data annotations, leading to three conclusions: (1) there is an effective exchange of metabolites among the two microalgae, (2) S. marinoi is most affected by this co-culture setup, and (3) there is a significant increase in the number of differentially expressed ORFs for both *S. marinoi* and *P. parvum*, which suggests adaptation. This work signifies that much remains undiscovered in our understanding of the marine biosphere, and the microalgae metabolic interactions should be further explored.

## 4 Methodology

### 4.1 Inoculation preparation

We cultivated two microalgal species in this co-culture experiment, *Skeletonema marinoi* (RCC75) and *Prymnesium parvum* (UTEX2797). The co-culture experiment was conducted using 13 co-culture chambers (4x treatment/control (2) + 1 media blank), each with a capacity of 600 mL (15, 16). The co-culture chambers were separated into two chambers by a permeable membrane, the Durapore® Membrane Filter, with a pore size of 0.22 µm. The control groups of the two microalgae were represented by mono-cultures of each microalga grown as single species in a co-culture chamber (*S. marinoi* / *S. marinoi* and *P. parvum* / *P.parvum*), while the treatment groups were grown as co-cultures (*S. marinoi* / *P.parvum and P.parvum* / *S. marinoi*). 500 mL of Artificial Sea Water (ASW) media was added to the co-culture chambers, assuming an error of ±10 mL. To prepare the inoculum, exponentially growing diatom cultures were pooled into 2L bottles, and a sample was taken and measured for chlorophyll *a* fluorescence to normalise photosynthetic potential using a 96-well plate. Dilution was done to ensure the diatoms had roughly the same chlorophyll *a* fluorescence. A 50 mL inoculation of dense diatom culture, one with *S. marinoi* (NCBI-Taxonomy ID: 267567), and the other inoculation with *P. parvum* (NCBI-Taxonomy ID: 97485), were added to each half of the co-culture chambers representing the treatment group (i.e. one chamber with *S. marinoi*, while other half with *P. parvum*). Both halves were inoculated with either *P. parvum* or *S. marinoi* to prepare mono-culture samples as control groups. A 1 mL sample of inoculation culture was preserved in a 1.5 mL Eppendorf tube with 1% Lugol’s solution. A sample of 1 mL was taken from each half chamber for chlorophyll *a* fluorescence and Lugol’s preservation for day 0. The co-cultures were moved to an 18°C chamber with cork rings for standing chambers and a 14/10 day/night cycle. The light intensity ranged from 47-63 µmol/m2/s, and the co-cultures were shaken daily to promote adequate diffusion of metabolites and randomise the positions.

### 4.2 Sample preparation

For sampling, 1 mL syringes were used with long syringe needles daily to extract diatom cultures from the randomly located chambers. A 96-well plate was used for chlorophyll *a* fluorescence measurement (200 µL), and a 1.5 – 2 mL Eppendorf tube for sample preservation (800 µL) using 1% Lugol’s preservation solution. Sampling was carried out every other day, and the regimen was days 0 (inoculation), 2, 4, 6, and 8, with cultures sampled right before extraction on day 8. After sampling, 50 mL of shaken culture was collected, discarding most of the filtrate. 2 mL of filtrate was stored for nutrient analysis in a freezer at -80°C. 5 mL Eppendorf tube filters were labelled for transcriptomics. Once dry, the filter was replaced, and 50 mL diatom culture was added to preserve two technical replicates for each biological replicate. After transcriptomics filters were collected, a new filter was added to the funnels and attached the funnels using tubing to a preconditioned SPE cartridge on the manifold. 50 mL of each culture for metabolomics extraction was filtered. The filters were saved in a 5 mL Eppendorf tube for endometabolomic extraction. We restarted the procedure with a new set of co-culture chambers and repeated it until it was done. We saved samples (argon sparged) in a -80°C freezer in a labelled box. Figure 4 illustrates the different co-culture chambers with control and treatment groups and the omics approaches applied to the samples.

**Figure 4:**
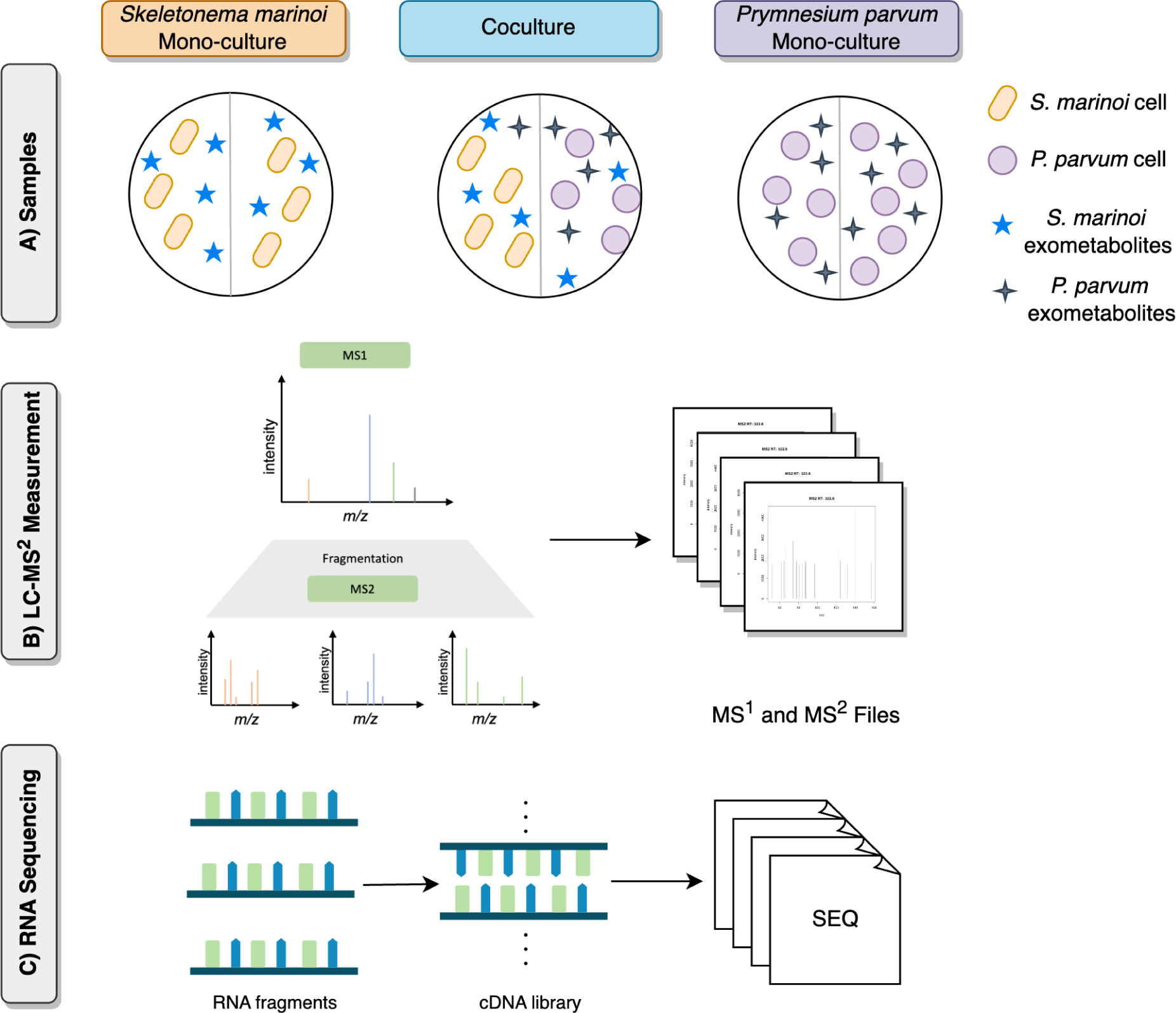
Experimental setup for metabolomics and transcriptomics analysis of co-culture of *Skeletonema marinoi* and *Prymnesium parvum*. A) The control groups were *S. marinoi* mono-culture and *P. parvum* mono-culture, while the treatment group was *S. marinoi* in the presence of *P. parvum* and *P. parvum* in the presence of *S. marinoi*. B) From the extracts of these groups, we acquired metabolomics data using LC-MS^2^, and C) RNA sequencing data.

### 4.3 Metabolomics

#### 4.3.1 Data Acquisition

Ultra High-Performance Liquid Chromatography (UHPLC) coupled with High-Resolution Mass Spectrometry (HR-MS) was carried out with Thermo (Bremen, Germany) UltiMate HPG-3400 RS binary pump, WPS-3000 autosampler, and THERMO QExactive plus orbitrap Mass Spectrometer coupled to a heated electrospray source (HESI). The separation was run on a Thermo 100mm, C18 RP (100 × 2.1 mm; 2.6 µm) column with a gradient of 100% A (Water+2% acetonitrile + 0.1% formic acid) to 100% B1 (acetonitrile) in 8 min and keeping 100%B1 for 3 min. All biological replicates were profiled via electrospray ionisation (ESI) in both positive and negative modes with a resolution of 70,000 and a mass range of 80-1200 *m*/*z*. A pooled quality control sample, made from 20 µL of each normalised (to cell count) biological replicate, was profiled after every five biological replicates for retention time shift corrections via the Compound Discoverer software (Thermo Scientific, version 3.2). Media blank samples were also profiled via LC-MS to remove all metabolites originating from media components. The quality control samples were used to generate a master inclusion list of metabolites of interest derived from phytoplankton metabolism. This list included 1467 features and 758 features for the exometabolomic and endometabolomic samples, respectively. The master inclusion list was separated into smaller, 100-feature lists for ease of data collection. Each 100-feature list included features spanning the full retention time span of the analysis window. The quality control samples were then profiled with each inclusion list via data-dependent acquisition mode (DDA) to generate fragmentation patterns for each parent ion included in the master inclusion list. The DDA analysis was also conducted in both positive and negative modes, using a three-stepped normalised collision energy of 15, 30, and 45. Figure 4 illustrates the metabolomics data acquired from this experimental setup.

#### 4.3.2 Data Preprocessing and Statistical Analysis

The MS^1^ processing workflow was developed with the LC-MS data generated using the mono-culture and co-culture samples. An overview of the workflow is given in Figure 2. The LC-MS spectrometry data RAW files were converted to mzML open format processed by the GNPS “Conversion Drag and Drop” file converter (58) (accessible at https://gnps-quickstart.ucsd.edu/conversion). Pre-processing and analysis were done in R, based on the “Identifying ESsenTIal Molecular variables in Terrestrial Ecology” (iESTIMATE) repository (59). The processing was mostly performed with the packages XCMS, Spectra, and MSnbase (60). In the following processing steps, the parameters were optimised for four different conditions: endometabolome in positive and negative modes and exometabolome in positive and negative modes. The parameters for the peak picking algorithm were optimised using the R package Isotopologue Parameter Optimization (IPO) (61).

The first function in the workflow is the data_preparation function. This function generated a descriptive table containing metadata and phenotypic data, called the *phenodata* table. The data were subset to an instrument-specific retention time range of 0-700 seconds and MS level = 1. Chromatographic peak picking was performed by the peak_detection function using the centWave algorithm (62) from the XCMS package (findChromPeaks). The following parameters were used for the function grouping_1, which used groupChromPeaks for grouping of peaks: minFraction = 0.7, bw = 2.5 (standard deviation of smoothing kernel). One resulting peak group was termed as a feature, representing a metabolite in the samples. Retention time correction was performed in the rt_correction function by adjustRtime with the peakGroups method. The following parameters were used: minFraction = 0.7, smooth = loess, span = 0.5, and family = gaussian. With the function grouping_2 a second grouping was performed with the same parameters as before. A feature table was created in the feature_extraction function using the featureValues function. Intensities of the feature table were transformed logarithmically, and missing values were imputed to 0 in the cfeature_transformation function. The lower end of the normal distribution of the feature data was used as an intensity cutoff for the creation of a presence/absence table, with a 5% cutoff in the binary_list_creation function. The workflow with documentation is available at https://github.com/zmahnoor14/MAW/tree/main/co-culture. The data is forwarded 1) to link MS^1^ precursor masses to their fragmentation MS^2^ spectra and 2) for statistical analysis.

For univariate statistical analysis, the chlorophyll *a* fluorescence was measured through 0 to 8 days of cell growth in mono-culture and co-culture samples. Two-way ANOVA with replicates was performed to verify if there was a significant change in the chlorophyll *a* fluorescence between mono-culture and co-culture conditions for each species separately. As a posthoc test, we also performed Welch t-test (two-tailed t-test with a sample of unequal variance). For the metabolomics data, diversity analysis on feature level was performed using the presence/absence matrix, measuring the metabolite richness by calculating the Shannon diversity index (H’) for each sample origin using R package vegan’s diversity function with index = Shannon. Multivariate statistical analysis was performed on the feature tables from the MS^1^ workflow to elucidate differentially abundant features. Principal component analysis (PCA) was performed using the prcomp function (63). Partial least square (PLS) analysis was performed using the f.select_features_pls function from the iESTIMATE. The parameters for model construction were principal components = 5, tune length = 10, and quantile threshold = 0.95.

#### 4.3.3 Suspect List Preparation and Metabolome Annotation

For metabolite annotation, we generated, based on literature and public databases search, two separate lists of metabolites known to be produced by both microalgae, termed suspect lists. In a previous study, the suspect list for *Skeletonema* spp was published (51). For the present study, we assembled a suspect list for *Prymnesium parvum* as well, using different databases: compound, reaction, pathway, and enzyme databases such as Kyoto Encyclopedia of Genes and Genomes (KEGG) (64–66), Comprehensive Marine Natural Products Database (CMNPD) (67), Chemical Entities of Biological Interest (ChEBI) (68), PubChem (69, 70), Universal Protein Resource (UniProt) (71), BRaunschweig ENzyme DAtabase (BRENDA) (72), MetaboLights (73, 74), MetaCyc (75, 76), and different publications on *P. parvum* metabolites (77–82). Suspect list was curated with RDKit (83), PubChemPy (84), and pybatchclassyfire (85). The protocol for curation is given in the methodology section of (51).

The chemical structure annotation was performed using the Metabolome Annotation Workflow (MAW) (22). All four conditions (endo_pos, endo_neg, exo_pos, and exo_neg) were combined into four .mzML input files for MAW. To perform dereplication, MAW-R (spectral database dereplication module) used GNPS (58), HMDB 5.0 (86, 87), and MassBank (88) (DOI: 10.5281/zenodo.7519270) (89), and MAW-SIRIUS (compound database dereplication module with SIRIUS5 (90), using COCONUT database (23). Different candidates from MAW-R and MAW-SIRIUS were analysed to select top candidates using MAW-Py. Chemical classes were assigned using ClassyFire (91)and CANOPUS (92). Finally, the putative candidate list obtained from MAW was screened for common compounds within the suspect lists of both species. The workflow was executed using docker containers with R and Python scripts (https://github.com/zmahnoor14/MAW) (93).

#### 4.3.4 Linking Annotations to Experimental Conditions

Following the MS^1^ preprocessing, statistical analysis, and MS^2^ annotation workflow, the MS^1^ features were linked to MS^2^ fragmentation spectra using inclusion lists to identify the chemical structures assigned to each differentially produced metabolite. Note that here the MS^1^ features are termed as metabolites. To accomplish this, a function named linking_with_inc_list was implemented, which searched for MS^2^ precursor masses [*m*/*z*] in the inclusion list within the specified [*m*/*z*] minimum and maximum range of the MS^1^ feature list. If the condition was met, the retention time range from the inclusion list was used to search for the retention time median value in the MS^1^ feature list. The corresponding feature ID was allocated to the MS^2^ precursor mass when the second condition was also fulfilled. In cases where retention time median values did not precisely align within the rt min-max range, this information was noted as a comment alongside the corresponding feature ID. After establishing the link between the MS^1^ feature ID and the MS^2^ precursor mass, the precursor masses were searched in the results obtained with MAW, allowing for the assignment of feature IDs to chemical structure annotations. This was accomplished using a function called link_ann_inc, which linked the inclusion list feature ID to each precursor mass in the MAW results, applying the same conditions for precursor mass and retention time ranges. Subsequently, the MAW results were provided with the corresponding feature IDs and conditions leading to a structure assignment to differentially produced metabolites (https://github.com/zmahnoor14/MAW/blob/main/co-culture/linking.R). The complete metabolomics workflow is illustrated in Figure 5.

**Figure 5:**
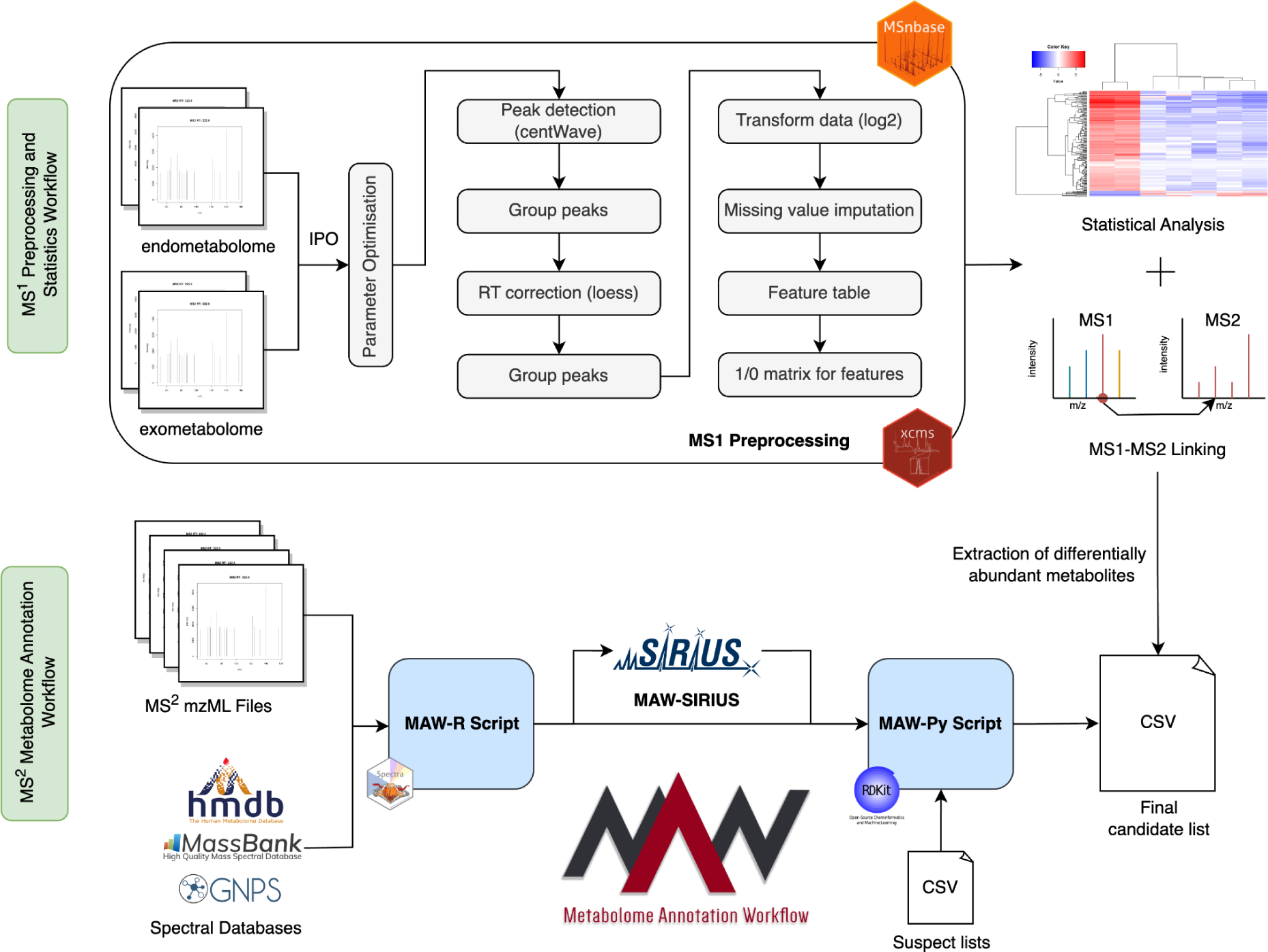
Metabolomics Workflow for the co-culture experiment. The first section demonstrates the MS^1^ preprocessing workflow, which starts with the .mzML files. These files go through the parameter optimisation. The optimised parameters are then used in the subsequent steps, such as peak detection, grouping, retention time correction, and re-grouping of peaks. The preprocessed data is then converted to a feature list with mass-to-charge ratios, retention time values, and intensities. These feature lists are subjected to further statistical analysis and are used to link the MS^1^ features to their corresponding MS^2^ fragmentation spectra. Separately, the MS^2^ Fragmentation spectra undergo Metabolome Annotation Workflow (MAW), which annotates chemical structures to the metabolic features using spectral databases and SIRIUS5. The final candidate list is searched in the suspect list from each origin organism to find previously identified metabolites in the new experimental setup dataset.

### 4.4 Transcriptomics

#### 4.4.1 Data acquisition

Total RNA was extracted using the Qiagen RNeasy Plant Mini kit and Plant RNA isolation aid kit following the manufacturer’s instructions (Qiagen, Hilden, Germany). RNA samples were quantified using Qubit 4.0 Fluorometer (Life Technologies, Carlsbad, CA, USA), and RNA integrity was checked with RNA Kit on Agilent Tapestation (Agilent Technologies, Palo Alto, CA, USA). The RNA sequencing library was prepared using the NEBNext Ultra II RNA Library Prep Kit for Illumina using the manufacturer’s instructions (New England Biolabs, Ipswich, MA, USA). Briefly, mRNAs were initially enriched with Oligod(T) beads. Enriched mRNAs were fragmented for 15 minutes at 94 °C. First-strand and second-strand cDNA were subsequently synthesized. cDNA fragments were end-repaired and adenylated at 3’ends, and universal adapters were ligated to cDNA fragments, followed by index addition and library enrichment by PCR with limited cycles. The sequencing library was validated on the Agilent TapeStation (Agilent Technologies, Palo Alto, CA, USA) and quantified by using Qubit 2.0 Fluorometer (ThermoFisher Scientific, Waltham, MA, USA) as well as by quantitative PCR (KAPA Biosystems, Wilmington, MA, USA). The sequencing libraries were multiplexed and clustered onto a flow cell. After clustering, the flow cell was loaded onto the Illumina HiSeq 4000 or equivalent instrument according to the manufacturer’s instructions. The samples were sequenced using a 2×150bp Paired-End (PE) configuration. The HiSeq Control Software (HCS) conducted image analysis and base calling. Raw sequence data (.bcl files) generated from Illumina HiSeq was converted into Fastq files and de-multiplexed using Illumina bcl2fastq 2.20 software. One mismatch was allowed for index sequence identification. Figure 4 illustrates the transcriptomics data acquired from this experimental setup.

#### 4.4.2 RNA seq Quality control and Transcriptome Assembly

Raw sequencing reads were assessed for quality using FastQC (version 0.11.9) (https://www.bioinformatics.babraham.ac.uk/projects/fastqc/). Adaptor trimming, quality filtering, and read preprocessing were performed using fastp (version 0.23.2) (94). In detail, 5’ and 3’ bases with a Phred quality score below 30 were cut, and reads with more than 1 N base, an average quality score below 30, or a length of fewer than 15 bases were removed.

Before assembly, the quality filtered reads were classified using Kraken2 (version 2.1.2) (27) using its standard index and including the genome assemblies of the diatoms *Thalassiosira pseudonana* (GCF_000149405.2, (95)) and *Skeletonema costatum* (GCA_018806925.1, (96)). All reads classified as viral, prokaryotic, and human were removed from the datasets. Trinity (version 2.13.2) (26) was used to assemble the transcriptome of the remaining read data, keeping only contigs with a minimal length of 200 nt. After assembly, Kraken2 was applied again to filter all assembled contigs not classified as originating from the clade Stramenopiles. BUSCO (version 5.4.5) (28) was applied to evaluate the assembly quality using the transcriptome mode and the stramenopiles_odb10 evaluation dataset.

#### 4.4.3 Transcriptome Annotation and Differential Gene Expression

Using the tool Augustus (29) (version 3.5.0), the putative open reading frames (ORFs) encoded by the assembled transcripts were annotated with the species option applied for *Skeletonema costatum* and the option to search on both strands. Using InterProScan (version 5.59_91.0) (34) and based on the databases Pfam (97), TIGRFAM (98), PANTHER (99), ProSiteProfiles (100), FunFam (101), the predicted protein sequences were further classified with their potential function, functional domains, as well as, associated GO terms and KEGG pathways. Processed reads were aligned to the assembled transcriptome using Hisat2 (version 2.2.1) (102) with standard parameters. Read counting was performed using featureCounts (version 2.0.1) (103), with the annotation described as above as a reference. Differential gene expression analysis was performed using DESeq2 (version 1.38.3) (104) stastical significance defined as FDR adjusted p-value < 0.05. The source code is available at https://github.com/Bioinformatics-Core-Facility-Jena/SE20220705_97.

### 4.5 Data Availability

The RAW and mzML metabolomics data are submitted and under curation on the MetaboLights (73, 74) repository with the study number MTBLS8313 (www.ebi.ac.uk/metabolights/MTBLS8313). The mzML files are also available on Zenodo with the following DOIs: (LC-MS^1^ 10.5281/zenodo.10143127; LC-MS^2^ 10.5281/zenodo.10143233 (105, 106)). All the metabolomics result files are available at Zenodo with DOI: 10.5281/zenodo.10143554 (107). The currently available versions of the GNPS, HMDB, and MassBank are also archived on Zenodo with the DOI: 10.5281/zenodo.7519270 (89). The suspect list for *Skeletonema* spp. is available on Zenodo with the DOI: 10.5281/zenodo.5772755 (108) and for *Prymneisum parvum* with the DOI: 10.5281/zenodo.7864506 (109). The script for the suspect list curation is available at https://github.com/zmahnoor14/MAW-Diatom/blob/main/suspectlist_curation.py (110), applied to both suspect lists. The code and scripts for MS^1^ processing, linking, and statistical analysis are available at https://github.com/zmahnoor14/MAW/tree/main/co-culture (111), while the MS^2^ processing and annotation were carried out using MAW v1.5.0 available on https://github.com/zmahnoor14/MAW (93). The transcriptomics sequencing data is available on the BioProject PRJNA1006530 (112). The fasta sequences for the proteins encoded by the predicted ORFs are also archived on Zenodo with the DOI: 10.5281/zenodo.10397384 (31). Code for transcriptome analysis is available on the GitHub repository: https://github.com/Bioinformatics-Core-Facility-Jena/SE20220705_97 (113).

## Supporting information

Supplementary Results 1

## CRediT Author Statement

Mahnoor Zulfiqar wrote the manuscript, worked on annotations and MS^1^ and MS^2^ linking, helped with the development of MS^1^ preprocessing and statistical analysis pipeline, enabled FAIRification of data, code and workflow, and curated the *P. parvum* suspect list.

Anne-Susan Abel developed the preprocessing and statistical analysis pipeline, analysed the results for metabolomics, wrote the relevant methodology sections, and revised the manuscript.

Emanuel Barth performed the whole transcriptomics analysis using his developed pipeline on the sequencing data that was received from GENEWIZ, Azenta, wrote the relevant methodology sections, and revised the manuscript.

Kristy Syhapanha prepared samples, measured the chlorophyll a fluorescence, acquired MS1 and MS2 data, wrote relevant methodology sections, and revised the manuscript.

Remington Xavier Poulin supervised Kristy Syhapanha for the experimental design, sample preparation, and MS^1^ and MS^2^ data acquisition, and revised the manuscript

Sassrika Nethmini Costa Warnakulasuriya Dehiwalage acquired the suspect list of *P. parvum* from different sources

Georg Pohnert obtained funding for the project, supervised the project, specifically the experimental design, metabolomics data acquisition, and metabolomics analysis, and revised the manuscript for important intellectual points.

Christoph Steinbeck obtained funding for the project, supervised the project specifically for metabolomics and cheminformatics analysis, and revised the draft for important intellectual points

Kristian Peters supervised the project, specifically for the development of MS^1^ preprocessing and statistical analysis pipeline, deployed IPO on an external server to obtain optimised MS^1^ parameters, and revised the draft for important intellectual points.

Maria Sorokina conceptualised and supervised the whole project and revised the draft for important intellectual points.

Authors would like to acknowledge Nico Ueberschaar (ORCID: 0000-0002-4192-490X) for providing the standardised MS^1^ data acquisition protocol.

