## Supplementary Table 1 for "Analysis of Metabolomics and Transcriptomics Data to Assess Interactions in Microalgal Co-culture of *Skeletonema marinoi* and *Prymnesium parvum*"

Supplementary Table 1. Chlorophyll *a* fluorescence measurement in relative fluorescence units (RFU) for 8 days (d0-d8) in *S. marinoi* mono-culture (Sm control), *S. marinoi* co-culture (Sm co-culture), *P. parvum* mono-culture (Pp mono-culture), and *P. parvum* co-culture (Pp co-culture).

| Culture | Replicates | d0 | d2 | d4 | d6 | d8 |
|---|---|---|---|---|---|---|
| Sm mono-culture | r1 | 0.0988 | 0.2443 | 0.6976 | 1.1495 | 2.3438 |
| | r2 | 0.0708 | 0.2042 | 0.7805 | 1.2755 | 2.1737 |
| | r3 | 0.0830 | 0.2215 | 0.6733 | 1.2019 | 2.2858 |
| | r4 | 0.0577 | 0.2260 | 0.6656 | 1.4163 | 2.0770 |
| | r5 | 0.0830 | 0.2338 | 0.6351 | 1.2091 | 2.2419 |
| | r6 | 0.0589 | 0.2126 | 0.6183 | 1.2664 | 2.5691 |
| | r7 | 0.0795 | 0.2708 | 0.7472 | 1.2030 | 2.0671 |
| | r8 | 0.0840 | 0.2118 | 0.6363 | 1.2156 | 2.0812 |
| Sm co-culture | r1 | 0.0901 | 0.3254 | 0.5406 | 1.1137 | 1.6935 |
| | r2 | 0.0864 | 0.2457 | 0.4975 | 1.0347 | 1.4130 |
| | r3 | 0.1049 | 0.2102 | 0.5509 | 1.0367 | 1.9273 |
| | r4 | 0.1078 | 0.3009 | 0.6638 | 1.0197 | 1.5530 |
| Pp mono-culture | r1 | 0.0956 | 0.2193 | 0.3572 | 0.6797 | 0.8978 |
| | r2 | 0.0736 | 0.1884 | 0.3445 | 0.6998 | 0.8071 |
| | r3 | 0.0858 | 0.2019 | 0.3484 | 0.7013 | 0.8071 |
| | r4 | 0.0967 | 0.1813 | 0.3606 | 0.6963 | 0.9866 |
| | r5 | 0.0994 | 0.1906 | 0.3550 | 0.7119 | 0.8924 |
| | r6 | 0.0761 | 0.2244 | 0.3255 | 0.6973 | 0.9665 |
| | r7 | 0.0866 | 0.1861 | 0.3213 | 0.6356 | 0.9224 |
| | r8 | 0.0913 | 0.1799 | 0.2962 | 0.7196 | 0.8608 |
| Pp co-culture | r1 | 0.0919 | 0.2136 | 0.3870 | 0.6747 | 1.1451 |
| | r2 | 0.0845 | 0.1741 | 0.3316 | 0.6393 | 1.0614 |
| | r3 | 0.0860 | 0.1989 | 0.3319 | 0.7147 | 0.9810 |
| | r4 | 0.0835 | 0.1961 | 0.3154 | 0.6032 | 0.9791 |
