## Supplementary Table 2 for "Analysis of Metabolomics and Transcriptomics Data to Assess Interactions in Microalgal Co-culture of *Skeletonema marinoi* and *Prymnesium parvum*"

Supplementary Table 2. ANOVA analysis of variance for the species. The sample describes the overall difference in the data when comparing mono-culture to co-culture data. Days describe the difference in the data between the days of sample collection. The values determining a significant difference for the corresponding groups are marked in green. SS = sum of squares to represent the squared deviation from the mean, df = degrees of freedom, MS = mean square which is calculated by dividing (SS) by (df), representing the average amount of variation within each variation source, F = F-statistics to determine whether significant differences exist between groups and within groups, P-value = probability of the null hypothesis being accepted or rejected, F crit = critical F-value, the value from an F-distribution table that corresponds to the significance level of 0.05. Another indicator of significant variation among the samples for S. marinoi is the calculated F-value (F) being higher than the critical F-value (F crit). In contrast, the opposite holds for P. parvum.

| Source of Variation | SS | df | MS | F | P-value | F crit |
|---|---|---|---|---|---|---|
| S. marinoi mono-culture vs co-culture | | | | | | |
| Sample | 0.2574 | 1 | 0.2574 | 21.9032 | 5.74E-05 | 4.1709 |
| Days | 18.1273 | 4 | 4.5318 | 385.6088 | 2.54E-25 | 2.6897 |
| P. parvum mono-culture vs co-culture | | | | | | |
| Sample | 0.0052 | 1 | 0.0052 | 3.6002 | 0.0674 | 4.1709 |
| Days | 4.2965 | 4 | 1.0741 | 749.7051 | 1.37E-29 | 2.6896 |
