## Supplementary Figure 1 for "Analysis of Metabolomics and Transcriptomics Data to Assess Interactions in Microalgal Co-culture of *Skeletonema marinoi* and *Prymnesium parvum*"

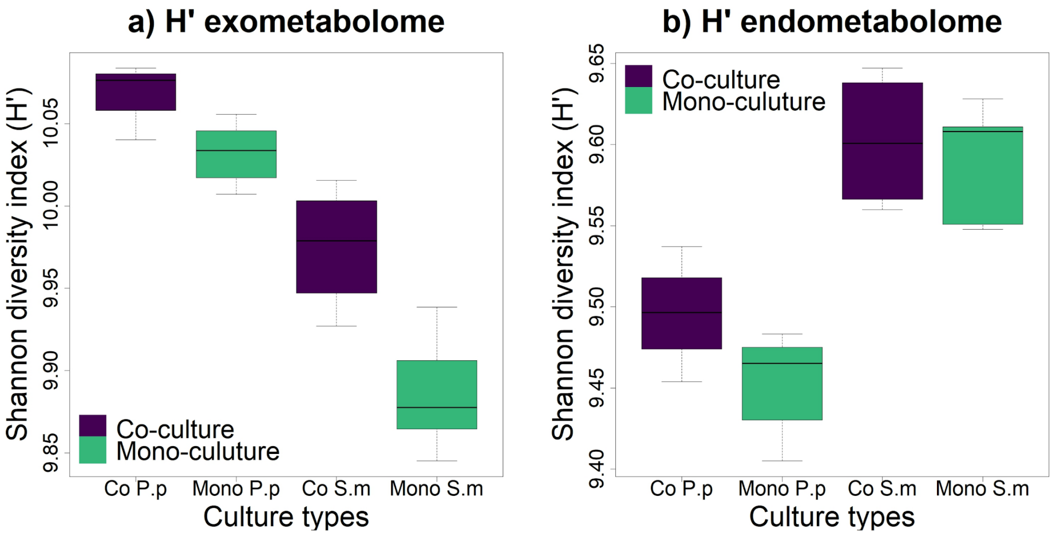


Supplementary Figure 1. Shannon Diversity index H’ for exometabolome and endometabolome samples, with co-culture samples coloured purple and mono-culture samples coloured green. a) exometabolome feature diversity with the left two samples showing *P. parvum* and the right samples *S. marinoi*. b) endometabolome feature diversity with *P. parvum* on the left and *S. marinoi* on the right. Mono = mono-culture, Co = co-culture, P.p = *P. parvum*, S.m = *S. marinoi*. For exometabolome, P. parvum showed higher chemodiversity of spectral features, with H’ = 10.03 for mono-culture and H’ = 10.07 for co-culture, than S. marinoi, with H’ = 9.89 for mono-culture and H’ = 9.98 for co-culture. For endometabolome, S. marinoi showed higher diversity, with H’ = 9.59 for mono-culture and H’ = 9.60 for co-culture, as compared to P. parvum, which shows H’ = 9.45 for mono-culture and H’ = 9.50 for co-culture.
