## Supplementary Figure 2 for "Analysis of Metabolomics and Transcriptomics Data to Assess Interactions in Microalgal Co-culture of *Skeletonema marinoi* and *Prymnesium parvum*"

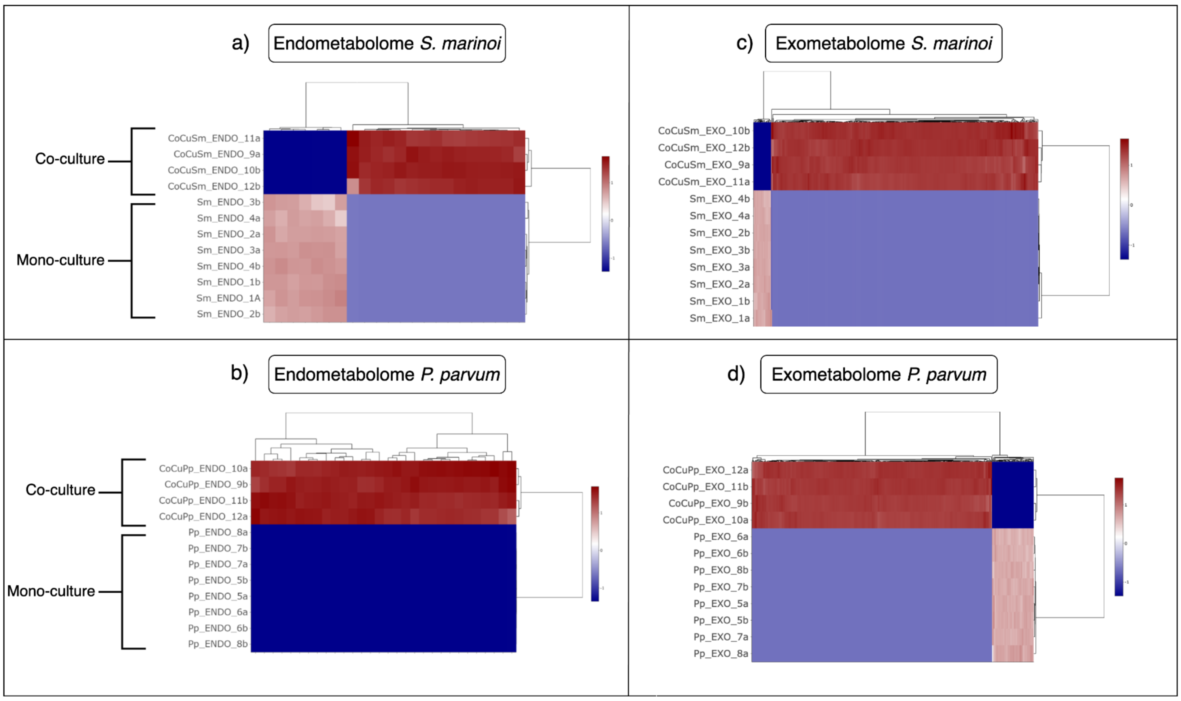


**Supplementary Figure 2:** Partial Least Square Analysis of endo and exometabolome of *S. marinoi* and *P. parvum*. The dendrogram clusters the conditions and features based on their intensities, with red indicating high intensity and blue representing low intensity or absence. a) For endometabolome, 22 differentially abundant features were selected for *S. marinoi*, out of which only seven were abundant in mono-culture. b) Conversely, all 30 differentially abundant features detected for *P. parvum* endometabolome were abundant in co-culture conditions. For exometabolome, a higher number of differentially abundant features were detected for both species, with more abundant features found in co-culture of *S. marinoi* and *P. parvum*. c) In total, the *S. marinoi* differentially abundant features were 324, out of which 304 features were abundant in co-culture. d) For *P. parvum,* the total number was 491, out of which 418 features were abundant in co-culture.
