## Supplementary Figure 3 for "Analysis of Metabolomics and Transcriptomics Data to Assess Interactions in Microalgal Co-culture of *Skeletonema marinoi* and *Prymnesium parvum*"

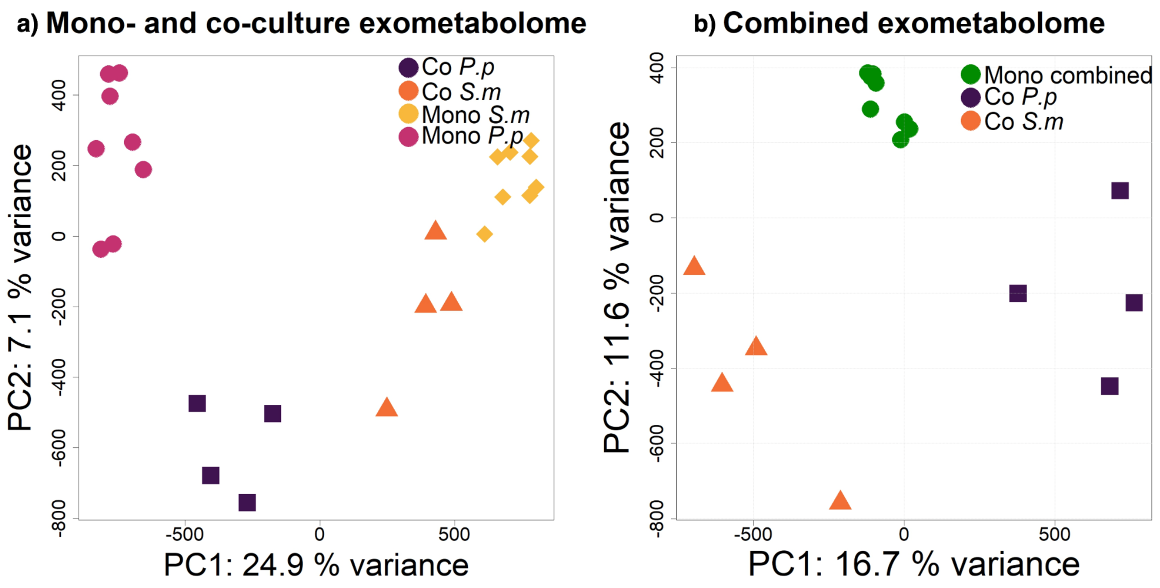


**Supplementary Figure 3:** Principal component analysis (PCA) for the observed effects on the exometabolome of co-cultures. a) Shows co-culture (orange for *S. marinoi* and purple for *P. parvum*) and mono-culture (yellow for *S. marinoi* and pink for *P. parvum*) samples with principal component (PC) 2, explaining 7.1 % of the variance, plotted against PC2, explaining 24.9 % of the variance. b) Shows combined mono-culture for the *S. marinoi* and *P. parvum* exometabolome in green, while the *S. marinoi* co-culture is shown in orange, and *P. parvum* co-culture is shown in purple, where PC 2 describes a variance of 11.6 %, and PC1 describes a variance of 16.7 %. The PCA in b) clearly distinguishes between the combined mono-cultures and individual co-cultures, indicating metabolic transformation occurring in the co-culture of both species.
