## Supplementary Results 1 for "Analysis of Metabolomics and Transcriptomics Data to Assess Interactions in Microalgal Co-culture of *Skeletonema marinoi* and *Prymnesium parvum*"

### Supplementary Results 1: Suspect lists from two marine microalgae

For this study, a suspect list of metabolites known to be produced by *P. parvum* was generated using publicly available databases and literature search. The suspect list for *P. parvum* consists of 222 primary and secondary metabolites, together with information on Formula, molecular mass, SMILES, InChI, IUPAC names, source, and ChemONT classification (1), which classifies chemical compounds based on their chemical structures into a hierarchical system. The suspect list and a sunburst plot can be accessed on Zenodo. We also used the published suspect list of *S. marinoi* to obtain the list of known metabolites from this diatom (2, 3), which consisted of 893 compounds from both *S. marinoi* and *S. costatum*.

The secondary metabolites detected from the *Skeletonema* spp. suspect list found in the endo- and exometabolome were: 7-mercaptoheptanoic acid, a medium-chain fatty acid, hexadeca-6,9,12-trienoic acid, which is a fatty acyl (4), and octatrienal which is a polyunsaturated aldehyde known to be produced by marine diatoms (5), and 8,11,14,17-eicosatetraenoic acid, all of which emphasise the diversity of fatty acids of *S. marinoi*. Toluene, a compound commonly associated with oceanic phytoplankton contributing to the remote marine atmosphere, was detected in the *S. marinoi* endometabolome (6). Finally, uteroverdine, which has a structure resembling biliverdin, was also found in the *S. marinoi* suspect list for the exometabolome. Lumichrome, being a blue-fluorescing compound formed by riboflavin photolysis, was present in *S. marinoi*'s exometabolome. Both *S. marinoi* and *P. parvum* shared the presence of fucoxanthin in their endometabolome, hinting at chlorophyll *a* production as part of the photosystems. Also shared between the two species were monopalmitin and monomyristin, glycerolipids previously identified in microalgae (7). *P. parvum*'s exometabolome showcased viburnitol, a cyclitol (8), which can have various roles, such as signal transduction, cell wall formation, and osmoregulation, and can act as antioxidants (9).
